# mRNA expression is co-regulated by non-nucleolar RNA polymerase I

**DOI:** 10.1101/2024.10.01.615958

**Authors:** Lucas M Carter, Ruyi Gong, Nicolas Acosta, Wing Shun Li, Emily Pujadas Liwag, Tiffany Kuo, Lam Minh Uyen Phan, Kyle MacQuarrie, Sui Huang, Masato T. Kanemaki, Luay Almassalha, Vadim Backman

## Abstract

The relationship between gene transcription and chromatin organization is an area of active study. Due to its role in mRNA synthesis, many studies have focused on the regulaton of RNA polymerase II (Pol-II) function by supranucleosomal structure and vice-versa. In contrast, there is little work on the function of RNA polymerase I (Pol-I) in non-nucleolar chromatin. Prior work has shown that Pol-I engages with components of Pol-II on rDNA, but it’s role in global transcription and chromatin structure beyond the nucleolus has largely been ignored. By pairing auxin-inducible degron technology with nanoscopic imaging, RNA-Seq, and Hi-C, we found that Pol-I and Pol-II co-regulate conformationally defined chromatin domains and mRNA synthesis. Mechanistically, Pol-I maintains the positioning of intronic and intergenic chromatin within domains for the proper expression of exon elements. Consequently, Pol-I loss disrupts genome connectivity, *in situ* chromatin domains, and the expression of mRNA, genome-wide.

## Introduction

The irregular 10-nm nucleosome “beads-on-a-string” chromatin heteropolymer is the fundamental organizational unit of the mammalian genome. Above this scale, chromatin organizes into a variety of topological and structural features^1^. Topologically associated domains (TADs), chromatin loops, and A/B compartments, observed using chromatin conformation capture (3-C, 5-C, Hi-C) and multiplexed fluorescent in situ hybridization (merFISH, ORCA, etc) modalities, are found in all eukaryotic organisms, indicative of a robust and complex genome connectivity framework. The coupling of these connectivity features with the space filling properties of chromatin *in situ* is a complex undertaking due to the various limitations of Hi-C, super resolution microscopy, and electron microscopy. For example, super-resolution imaging modalities, like single molecule localization microscopy (SMLM) and structured illumination microscopy (SIM), can provide subdiffractional molecular-specific insight into chromatin structure, but the reconstructed structures depend on a ground-truth model of chromatin that is provided by electron microscopy^2^. Chromatin specific electron microscopy (ChromEM) has emerged to provide information about the ground-truth of the polymeric folding of the genome in 3-D space but lacks specificity at the level of individual genes or post-translational modifications^3^.

Understandably, due to the challenge of bridging the connective and physical structures of the genome, the structure-function relationship between transcription and supranucleosomal 3-D structure remains an area of open debate. For example, independently degrading all three polymerases produced minimal changes in TADs or compartments in high-throughput chromatin conformation capture (Hi-C)^4^. Likewise, global degradation of cohesin or CTCF, the regulators of TADs, does not lead to wide-scale disruption of transcription^5,6^, but nascent transcription is disrupted in the absence of either^7^. Although Pol-II depletion has little impact on TADs and compartments in interphase cells, Pol-II is necessary to reestablish contact features following the exit from mitosis^8^. In contrast to these studies leveraging Hi-C, imaging studies using transcriptional inhibitors have demonstrated profound changes in chromatin structure *in* situ. Specifically, chromatin compacts globally in cells treated with transcriptional inhibitors across modalities^9,10^. Relatedly, active Pol-II associates *peripherally* with high density conformationally defined chromatin domains with heterochromatic cores while disruption of transcription leads to domain swelling^3,10,11^. Likewise, modulation of *in-situ* chromatin domains can result in a transformation of ensemble gene expression that impacts cell development, chemoresistance, and malignancy^12–14^.

A handful of questions emerge from these contrasting findings regarding the genome structure-function relationship: 1) are interphase supranucleosomal chromatin domains organized a priori as an emergent property of the system or does the act of transcription generate *in situ* 3-D structure 2) How does chromatin structure regulate mRNA transcription? And 3) Does Pol-II act independently or does it coordinate mRNA transcription with other polymerases? To interrogate the relationship between transcription and structure, we explored the consequence of full transcriptional inhibition for all three polymerases using pharmological inhibition on 3-D genome organization. We then examined the individual contributions of Pol-I and Pol-II to supranucleosomal chromatin domain organization and subsequent regulation of mRNA expression. To this end, we took advantage of an AID2 degron line in the human colorectal carcinoma 116 line (HCT116) targeting the largest subunit of Pol-II^15,16^, POLR2A, and an original AID line targeting the largest subunit of Pol-I, POLR1A, in HCT116 cells^17^ for precise temporal control and rapid depletion of each target protein^15,18^. Using these lines, we compared the role of each polymerase in maintaining chromatin connectivity with Hi-C and conserving the *in situ* physical structure of chromatin domains with chromatin electron scanning transmission microscopy (ChromSTEM), partial wave spectroscopy (PWS), and multiplexed single molecule localization microscopy (SMLM). Finally, we examined the loss of either polymerase on total transcription using genome-wide expression analysis. Together, we identified an unexpected co-regulatory framework for both polymerases in which Pol-I activity constrains chromatin domains and the transcription of intronic DNA, while the loss of Pol-II results in aberrant 3’ transcriptional readthrough. In summary, we demonstrate a novel extranucleolar role for Pol-I function in gene transcription and 3D chromatin structure through maintaining chromatin domain integrity and connectivity.

## Results

### Inhibiting transcription globally disrupts in situ domains

Chromatin inhabits a dense physical nuclear nanoenvironment composed of regulatory proteins, ions, polymerases, RNA, and transcription factors. Above the level of individual nucleosomes, supranucleosomal chromatin domains range in physical size from 50 nm to 300 nm and range between ∼ 50 KB to 350 KB in one dimensional size. Domain behavior follows the physical principles of polymeric folding and display radially decreasing mass-density from the center to the periphery. These properties depend on the free energy of chromatin-chromatin interactions, chromatin-solvent interactions, enzymatically driven loop formation, and physical constraints within the nucleus^2,3,11,19–21^. Crucially, multiple lines of evidence suggest that chromatin domains are not simply the physical manifestation of TADs: they are (1) largely stable following RAD21 depletion, (2) directly regulate ensemble gene transcription, and (3) depend on polymeric folding in a constrained nuclear volume. In contrast, TADs crucially depend on RAD21 and CTCF for function, are not necessary to maintain transcription, and are not impacted by changes in nuclear volume^5,22^.

The physical properties of chromatin domains include fractal dimension (*D*), density (the chromatin volume concentration, CVC), and domain radius (r). Chromatin domains *in situ* fold based on the power-law polymeric properties of the genome and can be quantified by how mass scales with the space the chromatin polymer occupies, *M* ∝ *r*^*D*^, where *D* is the scaling coefficient. Inversely proportional to the space-filling properties of chromatin, the connectivity of the chromatin polymer in individual cells will decay as a function of genomic distance by the scaling exponent, *S*, that is also observed in the ensemble on Hi-C^3^. As we have previously shown, in population measurements *S* and *D* are inversely related, where 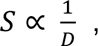 with biologically relevant *D* typically observed between 2 and 3^3,19^. Despite the reciprocal relationship between *S* and *D,* topological features such as TADs, loops, and A/B compartments do not behave consistently with the properties of chromatin domains^23^.

To untangle the interdependent relationship between domain structure and function in mammalian cells, we began by inhibiting transcription of all three polymerases with Actinomycin D (Act D), a potent chemotherapeutic agent, to study the consequence of complete inhibition of RNA synthesis on *in situ* physical structure. At nanomolar concentrations, Act D inhibits transcription of Pol-I rDNA repeats within the nucleolus, an effect due to the high rate of rRNA synthesis relative to mRNA^24^. At high concentrations (> 1 µg/mL), Act D intercalates DNA with a preference for GC-rich regions and completely disrupts transcription of all three polymerases^25,26^. We utilized *ActD* treatment at concentrations to inhibit the activity of all three RNA polymerases (> 1 µg/mL) for 1 hour and evaluated the effect on chromatin structure utilizing the previously described Nano-Chia platform as follows: live-cell Partial Wave Spectroscopy (PWS) nanoscopy, ChromSTEM tomography, and multi-plexed SMLM^3^.

PWS is a label-free live cell imaging technique that uses the interference in backscattered light from macromolecular structures such as chromatin to survey intracellular structure below the diffraction limit of light. Although individual domains are not directly resolved, PWS nanoscopy quantifies the proportion of chromatin organized into domains, *D*_*nucleus*_, which is proportional to the weighted average of the fractal dimension values of individual domains, 〈*D*_*domain*_〉, and the volume fraction of the chromatin domains. *D*_*nucleus*_ is also proportional to the fractional moving mass, FMM, defined as the product of the average mass of chromatin clusters that move coherently together per volume fraction of mobile chromatin, and the intranuclear effective diffusion rate of chromatin^27–29^. Consequently, increased *D*_*nucleus*_ represents a higher likelihood of chromatin organizing efficiently into domains and vice versa. Likewise, increased FMM represents coherent polynucleosome movement, consistent with an increased likelihood of domain formation events. Higher FMM might be due to having a greater volume fraction of mobile vs immobile chromatin or larger chromatin clusters moving as a single unit. The diffusion rate, better understood as the rate of change in chromatin mass-density, can be extracted from the temporal autocorrelation function decay curve by collecting the PWS signal over time at a single wavelength (550 nm). Diffusion provides a useful proxy for measuring the rate of macromolecular motion where smaller chromatin clusters typically correspond to a higher effective diffusion coefficient.

We first confirmed that transcription was fully inhibited in ActD-treated cells by labelling nascent RNA with 1 mM 5-Ethynyl-uridine for 1 hour following treatment with ActD for 1 hour and 6 hours at 5 µg/mL (**SI 1a,c**). Simultanenous labeling with DAPI followed by Coefficient of Variation analysis revealed compacted chromatin visible using widefield fluorescent microscopy (**SI 1b**). Upon ActD mediated inhibition, we observed an acute drop in packing scaling, *D_nucleus,_* and FMM in as little as 1 hour, consistent with the decreased likelihood of chromatin organizing into *in situ* domains throughout the nucleus (**Fig. 1a-d**). To confirm that this is a generalized phenomenon, we performed similar experiments in three additional cell line models with comparable results (**SI 1d**).

**Figure 1.**
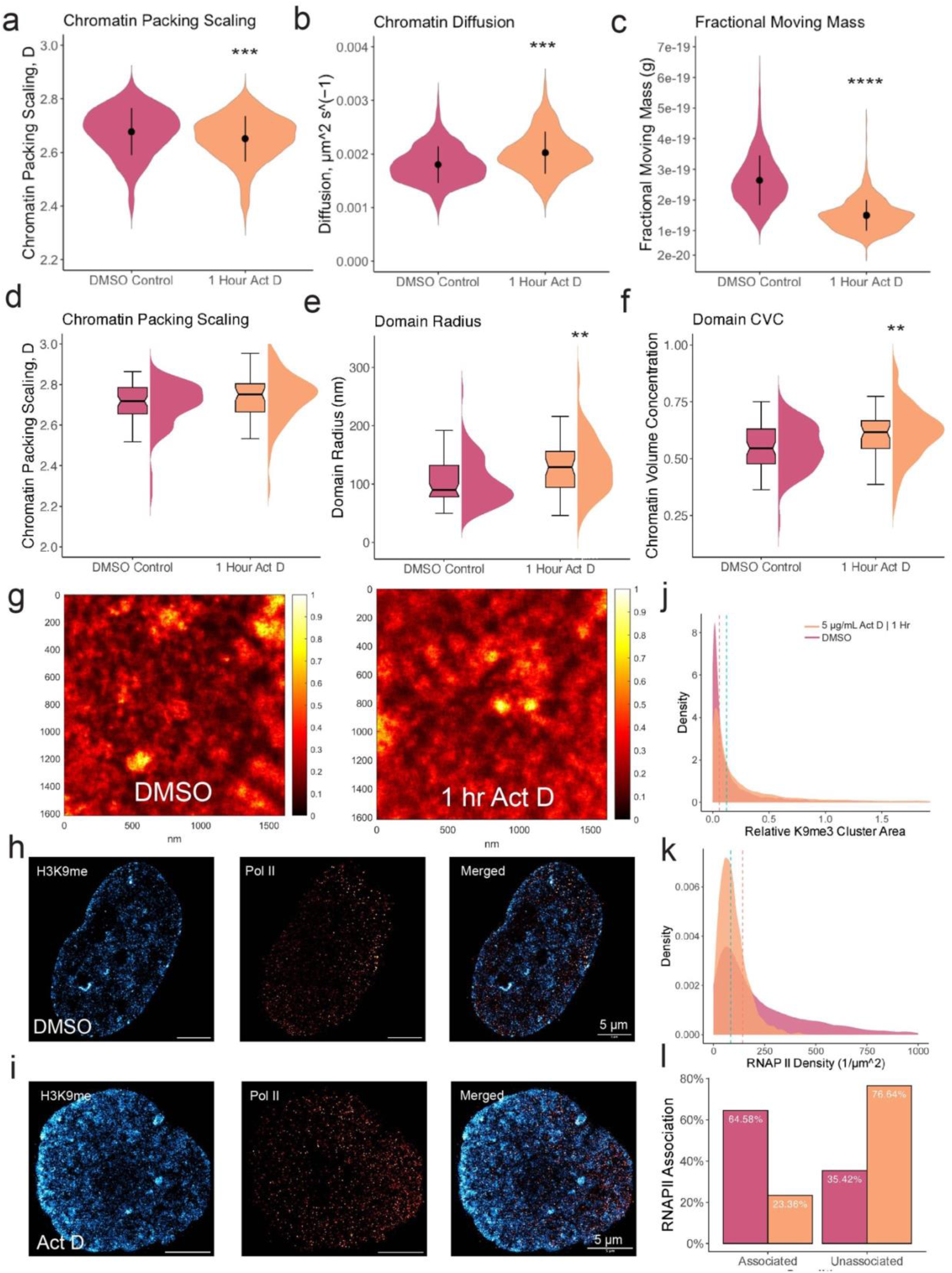
Chromatin domain analysis following transcriptional inhibition of HCT116 cells using Actinomycin D treated (5 µg/mL) or treatment with DMSO. (**A-C**) PWS following treatment: average nuclear packing scaling, diffusion, and fractional moving mass of chromatin domain nuclear average, respectively. (**D-F**) ChromSTEM following treatment: packing scaling, domain radius, and chromatin volume concentration of individual domains, respectively. (**G**) ChromSTEM tomograms showing untreated chromatin state or domain swelling following treatment. (**H-I**) Representative images of SMLM for labelled H3K9me3 (blue) and labelled active POLR2A (red). (**J-K**) Density distribution of H3K9me3 and POLR2A in SMLM analysis. (**I**) Association analysis of POLR2A with H3K9me3 clusters before and after treatment.

To understand the effect of transcriptional inhibition on individual domains directly, we analyzed ChromSTEM tomography of cells treated with ActD compared to untreated controls. Although ChromSTEM lacks molecular specificity, the resolved structures on ChromSTEM represent the geometric ground-truth of the genome *in situ* where the contrast is directly proportional to chromatin density with a resolution less than 5 nm. On ChromSTEM, we observed a loss of domains overall. Unexpectedly, we also observed that the size, density, and fractal dimension all increased for the remaining domains (+26.15 nm, +0.064 CVC, +0.028 D) on average (**Fig. 1d, e**). To understand how downregulation of chromatin domains translated to molecular modifications of nucleosomes and Pol-II localization, we utilized multiplexed SMLM, a super-resolution technique that resolves labelled molecules beyond the diffraction limit of light with precision down to 20 nm^30,31^. Using SMLM, we imaged the resulting transformation of transcriptionally active Pol-II (serine 2 phosphorylated, Pol-II Ps2) association with regions defined by constitutive heterochromatin (H3K9me3). Prior work had demonstrated both that Pol-II Ps2 localized to the transcriptionally active euchromatic periphery of chromatin domains and that constitutive heterochromatin comprised the dense interior of these domains^3,11^. Consistent with the findings on PWS nanoscopy and ChromSTEM tomography, the relative area occupied by H3K9me3-defined cores decreased by 0.064 clusters/µm^3^ following ActD treatment with a concurrent decrease in the density of Pol-II Ps2 loci (**Fig. 1j,k**). Visually, we observed a decoupling of POLR2A from heterochromatin (**Fig. 1h,i**). POLR2A, normally associated with heterochromatin cores (65%), disassociated from heterochromatin cores after Act D treatment (∼35%) (**Fig. 1l**). These results were consistent with prior work showing that POLR2A co-localized with nanoscopic domains and demonstrated that transcription was necessary for domain stability and maintenance in interphase cells^3,11^. Taken together, inhibiting transcription of all three polymerases produces an aggregation of chromatin into fewer but denser domains; Similar observations have been made using super resolution microscopy in which Act D generates an abrupt DNA-compaction phenotype across the nucleus^9,10^.

### Nuclear-wide gene transcription unexpectedly persists despite Pol-II depletion

Actinomycin D is a well-defined transcriptional disrupter of all three polymerases, with dose dependent selective inhibition of Pol-I, Pol-I and Pol-II, or all three polymerases simultaneously^32–34^. However, Act D has a complex mechanism of action rife with off-target effects that includes impacting topoisomerase function, supercoiled or quadraplex DNA, generating double strand breaks, and other non-specific interactions^35–37^. The lack of fidelity to a single mechanism makes interpreting Act D’s pleotropic phenotype challenging. Other transcriptional inhibitors, while more targeted than Act D, also disrupt non-transcriptional biological processes^26^. To overcome this limitation, we utilized two degron lines: one line in HCT116 cells targeting POLR1A using the original AID architecture and another in HCT116 cells targeting POLR2A with the improved AID2 system^15,17^. We chose to focus our attention on Pol-I and Pol-II because Pol-III accounts for only 10-15% of total cellular transcription and is less abundant than Pol-I or Pol-II^38–40^. The first iteration of the AID system, while providing valuable functional insight into essential proteins, was found to exhibit leakiness compared at baseline to its AID2 successor^41^. We compared our findings to both nongenetically modified wildtype HCT116 cells and DMSO-treated AID1 cells in our initial experiments for Pol-I. We found that the phenotype was similar between the wildtype and the degron line, but stronger in the wildtype line, likely attributable to inherent leakiness in the AID1 system. In later experiments, we only compared polymerase-degraded cells to wildtype HCT116 cells.

We verified the depletion of these proteins using widefield microscopy, flow cytometry and western blot analysis, confirming the complete degradation of POLR1A and POLR2A at 6 hours (**SI Fig 2a-b,d-e**). Additionally, widefield fluorescence microscopy confirmed that degron-tagged POLR1A, co-localized with nucleolin and UBF, formed characteristic polar caps upon 5 hours of auxin treatment (**SI Fig 2c**). We next sought to verify that the depletion of the respective polymerases resulted in the arrest of nucleolar rRNA (Pol-I) or mRNA gene transcription (Pol-II). ActD treatment completely inhibited transcription at 1 and 6 hours (**Fig 2a-b**), while POLR1A depletion resulted in the inhibition of nucleolar transcription. Surprisingly, although POLR2A depletion attentuated intra-nuclear transcription in the nucleoplasm, continued RNA synthesis was visually apparent in POLR2A-degraded cells (**Fig 2a-b**). To confirm that this was not an artifact of diffraction-limited resolution, we generated high quality SMLM imaging of nascent RNA labelled with 5-ethynyl-uridine (EU) in POLR1A and POLR2A degraded cells as previously described^41^ (**Fig 2c-e**). In both conditions, polymerase degradation increased the size of RNA clusters while RNA clusters became more diffuse. Surprisingly, RNA signal increased in POLR1A cells (**Fig 2f-h**). These results verified that loss of POLR1A leads to rRNA transcriptional abrogation. Again, we saw that POLR2A loss led to attenuated transcription in the non-nucleolar nucleoplasm but did not lead to full mRNA transcriptional inhibition.

**Figure 2.**
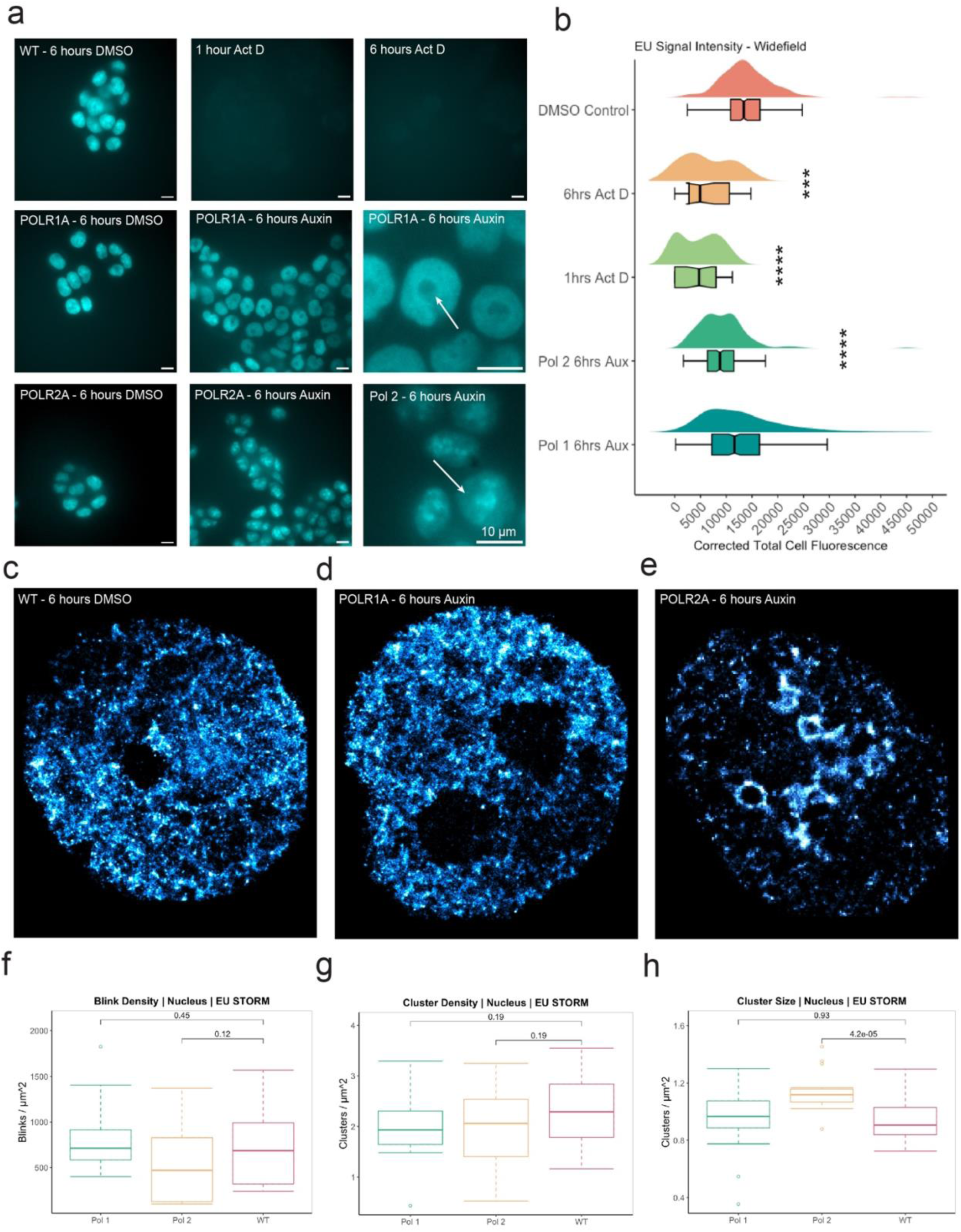
Degron lines (POLR2A-AID2 or POLR1A-AID1) were treated with auxin, DMSO, or Actinomycin D treated (5 µg/mL) for 6 hours and imaged following labeling of nascent RNA with 1 mM 5-ethynyl-uridine for 1 hour. (**A**) Representative image of each condition captured using widefield fluorescent microscopy. (**B**) Cell total corrected fluorescence of widefield fluorescent images for each condition. (**C-E**) Representative images from SMLM of EU-labeled RNA for each condition. (**F-H**) Quantification of SMLM distributions for each condition: fluorophore blink density, DBscan cluster density, and DBscan cluster size.

To verify that this unexpected result was not due to the transit of rRNA from the nucleolus, we performed total RNA-Seq, with rRNA depletion prior to sequencing, for ActD treated cells, POLR1A-degraded cells, POLR2A-degraded cells, and wild type HCT116 cells. Using a liberal cutoff (padj 0.05, abs(LFC) > 0.58), we observed broad suppression of gene transcription with ActD (**Fig 3a**) (**Fig SI 4a**). Consistent with our nascent RNA SMLM results, depletion of POLR2A triggered upregulation of upwards of ∼5500 low significance genes but only ∼2000 genes were downregulated (**Fig 3b**). Many of the genes that were upregulated in POLR2A degraded cells had low basal expression initially and experienced minor increases in expression (**Fig SI 4c**). These findings were reproducible and consistent across replicates, indicating that this was not an artifact of library preparation or from variable mRNA decays (**Fig 3k**). POLR2A degraded cells accounted for the majority of variance between all three conditions (**Fig SI 4d-e**).

**Figure 3.**
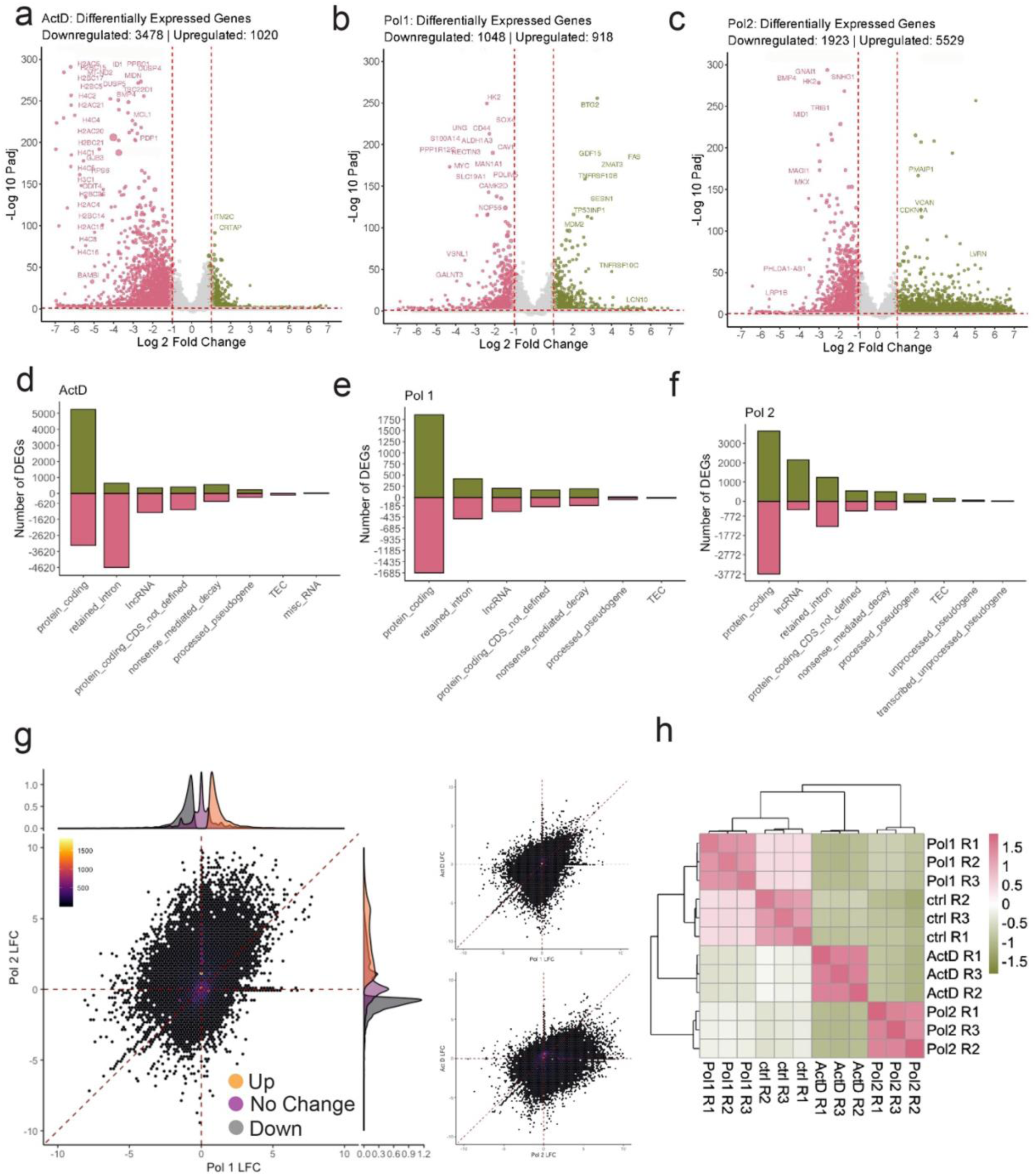
Degron lines (POLR2A-AID2 or POLR1A-AID1) were treated with auxin for 8 hours, DMSO for 8 hours, or Actinomycin D treated (5 µg/mL) for 6 hours and total RNA-seq for each condition was generated. (**A-C**) Volcano plots for each condition (P-adj < 0.05 | abs(LFC) > 1). (**D-F**) Number of DEGs for each type of transcript in each condition. (**G**) Scatterplot of LFC values for each gene in one condition compared to LFC values in another condition. Marginal distribution shows DEGs with same cutoffs as volcano plots. (**H**) Sample correlation.

Similar to POLR2A, degrading POLR1A led to broad differential expression throughout the genome. We observed 1048 genes that were downregulated and 918 genes that were upregulated (**Fig 3c**). Unlike ActD treatment or POLR2A depletion however, loss of POLR1A triggered even differential expression changes in both high and low basal expression protein coding genes. Comparison of POLR1A-degraded transcriptional patterns for genes in nucleolar associated domains (NADs), lamin associated domains (LADs), and intergenic chromatin also verified that this transformation was not confined to segments of the genome associated with the nucleolus (**SI Fig. 4d,e**). Comparing POLR1A to POLR2A depletion, we were surprised to discover that many genes that were downregulated in POLR1A depleted cells were reciprocally upregulated in POLR2A depleted cells and vice versa. Transcripts that underwent upregulation in either condition were generally downregulated in Act D treated cells (**Fig 3j**). We counted the annotations of transcripts that were differentially expressed in each condition. Independent of molecular function, Pol-I transcripts were equally up and downregulated across all biotypes, while Pol-II long non-coding RNAs, processed pseudogenes, and experimentally unconfirmed transcripts (TECs) were upregulated. Non-coding transcripts were primarily downregulated in ActD treated cells (**Fig 3g-i).**

**Figure 4.**
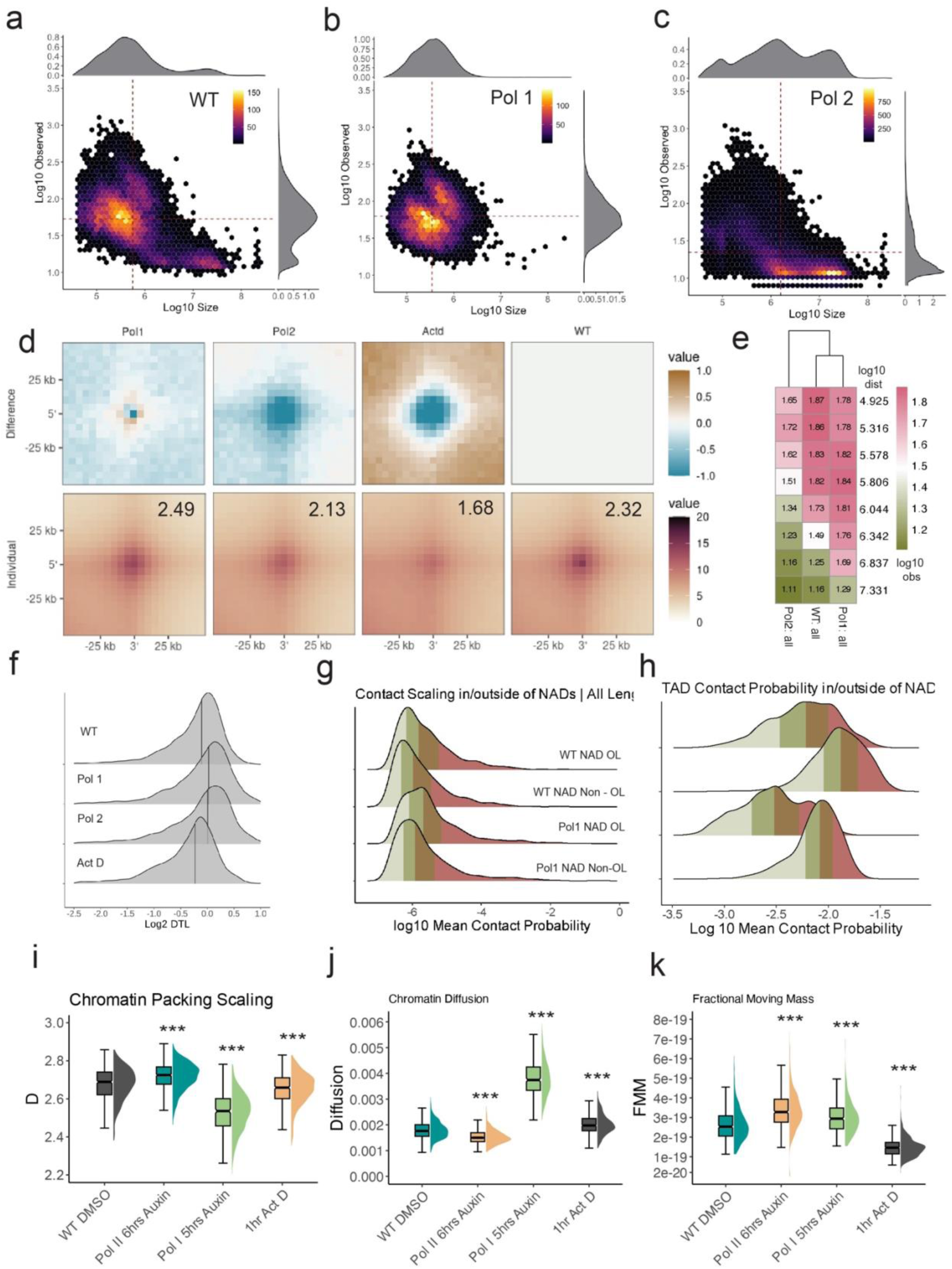
Degron lines (POLR2A-AID2 or POLR1A-AID1) were treated with auxin for 6 hours, DMSO for 6 hours, or Actinomycin D treated (5 µg/mL) for 1 hour and Hi-C was generated for each condition. (**A-C**) Scatterplot of log10 loop strength against log10 size for each loop. Loops called using HICCUPS. Heatmap shows loop density. (**D**) Top: difference plot showing change relative to WT. Bottom: Pileup plots of loop insulation strength for each. (**E**) Heatmap of loop strength binned by log10 loop distance. (**F**) Distribution of Distal-to-Local ratio values calculated in 10 kb bins for each condition. **(G)** Contact probability calculated in 10 kb bins for NAD regions and non-NAD regions. **(H)** Contact probability calculated within TADs for NAD regions and non-NAD regions. (**I-K**) PWS following treatment: average nuclear packing scaling, diffusion, and fractional moving mass of chromatin domain nuclear average, respectively.

### POLR1A and POLR2A co-regulate genome connectivity and in situ chromatin domains

Given these novel findings, we explored what processes could be facilitating the continued synthesis of mRNA in the absence of functional Pol-II and the global dysregulation of mRNA transcription following Pol-I disruption. We started by investigating the effect of POLR1A and POLR2A depletion on 1D chromatin connectivity via Hi-C and *in situ* structure using a combination of PWS nanoscopy, multiplexed SMLM, and Hi-C.

We generated biological Hi-C replicates for POLR1A degraded cells (871,928,451 contacts), POLR2A degraded cells (670,120,527 contacts), and ActD treated cells (502,274,008 contacts). These replicates were merged to maximize the resolution of each library, producing contact libraries with resolutions below 5000 BP (**SI 2a**). The individual experiments for each condition were assessed using a stratum-adjusted correlation coefficient and were found reproducible (**SI 2b)**^42^. Jiang et. al. previously reported limited impact on TADs and compartments upon degradation of each polymerase^4^. In agreement with previous studies using auxin-inducible degron technology or targeted drug inhibition, large-scale features of the genome such as TADs and A/B compartments do not undergo significant changes following degradation of either polymerase (**SI 5a-b,c-d**)^4,8,44^. Likewise, we observed little change in genome-wide relative contact probability at short and long distances (**SI 5g-h**). In contrast to Jiang et. al., POLR1A depletion resulted in a global loss of weak long-range loops above 1 Mbp, shedding 26.7% of all wildtype loops, while retaining more insulated short-range (< 1 Mbp) loops (**Fig 4a-e**). Conversely, POLR2A depletion resulted in an increase of many more weak entropic long-range loops (> 1 Mbp), gaining an additional 38,117 loops, while wildtype loops in POLR2A degraded cells were overall weaker (**Fig 4c-e**). One implication is that Pol-I may be primarily responsible for the formation of long-range looping events that were previously constrained by the activity of Pol-II whereas Pol-II may generate primarily short-range chromatin loops that were previously constrained by Pol-I.

In recent work, it was reported that heterochromatic NADs in mESCs were found on all chromosomes^44^ and nearly half overlapped with peripherally located LADs, making contact with non-nucleolar regions across the nucleus^45^. To assess the possibility that the impact of POLR1A is restricted to regions adjacent or tethered to the nucleolus, we exploited recently generated publicly available NAD data in HCT116 cells (**Table S1**). We segmented the human genome (hg38) into contact regions that spanned NADs or non-NADs into bins of 10 kb and measured the change in features outside of NADs in comparison to those restricted to within NADs. Consistent with the global change in gene transcription observed upon depletion of POLR1A, the transformation of chromatin loops, TADs, and contact scaling was conserved both within NADs and outside of NADs (**Fig. 4g-h**). We verified that these results were not unique to HCT-116 cells; we reanalyzed Hi-C data generated from mouse mESC degron lines where POLR1A and POLR2A were degraded upon auxin treatment^4^ along with publicly available nucleolar DNA adenine methyltransferase identification (DamID) data for NADs in mESCs^44^. Supporting our data, we found that NADs and non-NADs did not exhibit differential contact behavior at different length scales (**SI 5k).** These results underscore that the differential contact behavior observed following degradation of POLR1A is a global phenomenon related to supranucleosomal chromatin domain regulation, not a byproduct of nucleolar disruption.

To evaluate the contact behavior of topological features, we calculated the mean contact probability within these features. We found that mean contact probability was on average lower in POLR1A degraded cells in TADs (mean values here) (**Fig. SI 5l**). This behavior led us to ask if other contact regimes were perturbed. We calculated the Distal-to-Local (DTL) ratio of contacts between 50 kbp and 1 Mbp and contacts over 1 Mbp; we found that in both POLR1A and POLR2A degraded cells the total distribution of DTL values decreases, indicating shedding of local short range contacts in favor of long distance contacts (**Fig. 4f**)^46^. We then quantified the cis-trans ratio for POLR1A degraded cells. We observed that the rate of trans contacts decreased uniformly across all chromosomes by several percent each compared to wildtype cells (**Fig. SI 5j**).

We utilized PWS nanoscopy to investigate the consequence of the loss of each polymerase on chromatin in live cells. On live-cell nanoscopy, POLR1A and POLR2A depletion resulted in conjugated phenotypes with POLR2A depletion modestly increasing *D_nucleus_* and FMM. POLR1A depletion resulted in a pronounced drop in *D_nucleus_* and an increase in FMM. The change in FMM can be understood as an increase in the size of nucleosome clutches moving coherently (FMM) that indicates chromatin domain deterioration into large clutches of nucleosomes moving together (**Fig 4i,k**). We observed an acute decrease in the rate of movement of chromatin clutches in POLR2A degraded cells, but in POLR1A degraded cells the rate of movement increased (**Fig. 4j**). We also compared POLR1A and POLR2A degraded cells to DMSO treated cells of each respective degron line. The leaky degradation phenotype of the AID1 system in the POLR1A line is evident in this comparison, where baseline *D_nucleus_* is lower than wildtype cells (**Fig SI 8d-f).**

Using H3K9me3 labelling and SMLM, we characterized disruption of constitutive heterochromatin domain cores after full inhibition of Pol-I and Pol-II transcription. On SMLM, the depletion of POLR1A and POLR2A resulted in nuclear-wide reorganization of constitutive heterochromatin. In both conditions, we observed prominent swelling of heterochromatin at the nuclear periphery, but the swelling of heterochomatic lamina was particularly pronounced in Pol-I disrupted cells (**Fig 5a-c**). Inhibiting transcription of either polymerase resulted in heterochromatin cores spreading out across the nucleus: blink density (the concentration of K9) and cluster size both increased while the density of domain cores fell (**Fig 5d-f**). These results are consistent with earlier studies in which Pol-I transcriptional inhibition with nanomolar ActD disrupted both cis and trans contacts between heterochromatic NADs, nucleoli, and non-nucleolar chromatin^47,48^.

**Figure 5.**
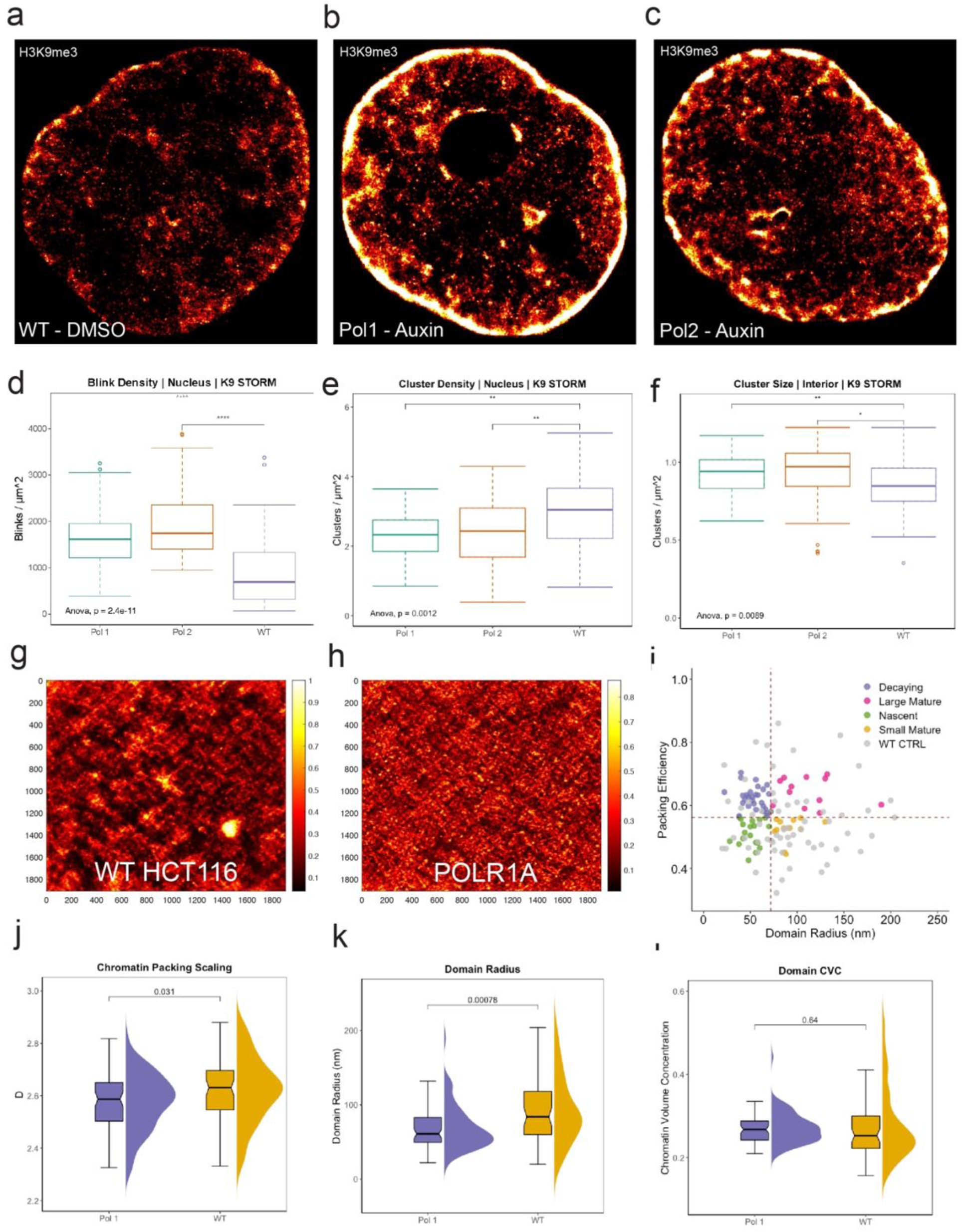
SMLM data was generated for degron lines (POLR2A-AID2 or POLR1A-AID1) were treated with auxin for 6 hours or DMSO for 6 hours, followed by ChromSTEM for the POLR1A-AID1 line under the same treatment conditions. (**A-C**) Representative images of H3K9me3-labeled SMLM. (**D-F**) Quantification of H3K9me3-conjugated SMLM distributions for each condition: fluorophore blink density, DBscan cluster density, and DBscan cluster size. (**G-H**) ChromSTEM tomograms showing untreated chromatin domains in untreated cells state or chromatin domain degradation in cells where POLR1A has been degraded. (**I**) Analysis of packing domains by size and packing efficiency to analyze domain properties. Nascent domains (low efficiency, small size), mature domains (high packing efficiency), and decaying domains (low efficiency, large size) represented by color. Lines represent median packing efficiency and median radius in control cells. (**J-L**) Distribution of individual domain properties observed on ChromSTEM: packing scaling, domain radius, and chromatin volume concentration of individual domains, respectively.

The transformation of heterochromatin alongside the decrease in *D_nucleus_* observed on PWS in POLR1A degraded cells led us to ask whether Pol-I had an extra-nucleolar role in chromatin domain regulation. While super resolution imaging is moleculary informative, we turned to chromSTEM tomography for ground truth information on what was happening to *in situ* individual chromatin domains upon inhibition of Pol-I transcription. In stark contrast to tomography of ActD treated cells, loss of Pol-I transcription precipitated disintegration of coherent chromatin domain organization (**Fig 5g-h, SI 6a**). The fractal dimension, 〈*D*_*domain*_〉, of individual domains measured on chromSTEM decreased from 2.62 to 2.58 on average, while domain radius decreased from 92 nm to 69 nm. Chromatin volume concentration increased on average across all individual domains (**Fig 5j-l**). We categorized the loss of domains following POLR1A degradation by their packing efficiency, *α*, and domain radius, R: 20% of decaying (large domains with low packing efficiency) or mature stable domains (large domains with high packing efficiency) were lost (**Fig 5j-l**)^23^. Together, these results suggest that Pol-I may function to stabilize mature chromatin domains outside the nucleolus and when lost, trigger domain degradation. In comparison, Pol-II appears to generate domains, potentially through driving enhancer-promoter and promoter-promoter transcriptional looping.

### Pol-I associates with intergenic and intronic non-nucleolar chromatin

Prior work has demonstrated that Pol-I can be located outside nucleoli^49^. Our earlier results indicated that POLR1A degradation led to nuclear disruption of chromatin domains across the nucleus. To rule out a purely nucleolar cause, we made use of multiplexed SMLM, publicly available ChIP-Seq data, and Cut&Tag to verify Pol-I association with extranucleolar chromatin. Using widefield fluorescent microscopy, we first confirmed that at low resolution, the bulk of Pol-I was visible within nucleolar structures, delineated by nucleolin staining (**Fig 6d**). Using SMLM, we imaged Pol-I simultaneously with constitutive heterochromatin (H3K9me3). While nucleolar regions were enriched in Pol-I, super resolution imaging revealed that Pol-I was also abundant within intergenic chromatin and near the lamina (**Fig 6a**). Within the non-nucleolar space, we observed that Pol-I colocalizes outside H3K9me3 heterochromatic cores at comparable levels of association and distances to that observed with actively transcribing Pol-II-Ps2 (**Fig 6a-c**). 65% to 81% of extranucleolar Pol-1 molecules were associated with heterochromatin cores. From reanalysis of ChIP-Seq data generated in mouse cells, Pol-I and Pol-II frequently colocalize with each other independent of the transcript type (miRNA, lncRNA, protein coding genes) both within NADs and outside of NADs (**Fig 6e**)^40^. Furthermore, Pol-I shows no preference for NADs or non-NADs. Quantitatively, POLR1A and POLR2A peaks frequently colocalized within 10kbp segments with a spearman correlation coefficient of 0.55. This contrasts with the likelihood of Pol-I or Pol-II random co-positioning within any genomic segment at 10kbp resolution (correlation coefficient < 0.1, **Fig 6f**). We verified the genome-wide binding of Pol-I in a human ChIP-seq dataset as well. POLR1A peaks were found widely distributed across all chromosomes in human mammary epithelial cells (HMEC) (**Fig 6g**).

**Figure 6.**
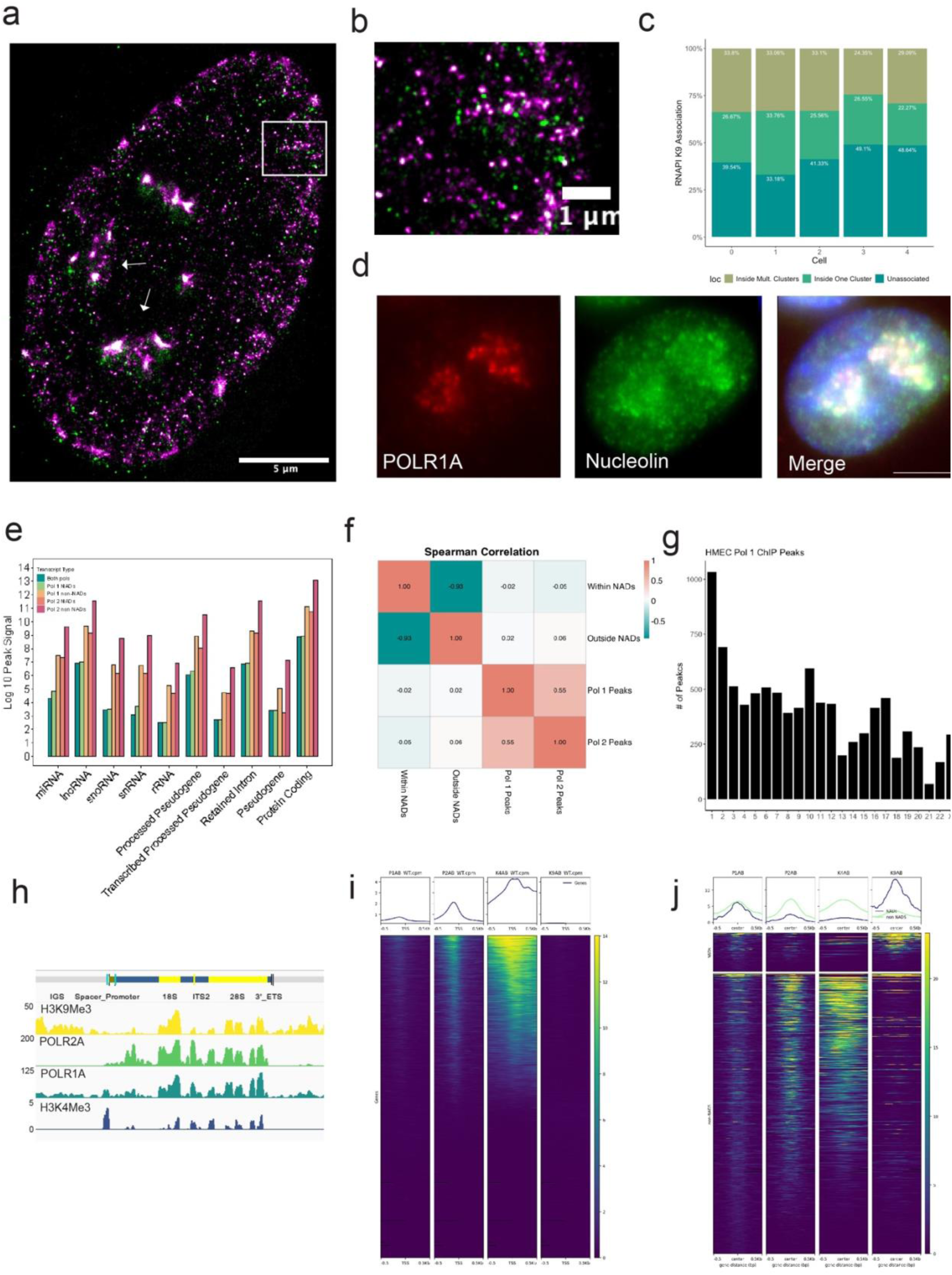
Pol-I is widely bound on non-nucleolar genomic elements. (**A**) SMLM image of labeled extrannucleolar POLR1A (green) and labeled H3K9me3 (magenta) denoting heterochromatic cores. (**B**) Magnified view of (**A**) showing POLR1A on the periphery of a heterochromatic core. (**C**) Association analysis of H3K9me3/POLR1A SMLM showing percentage of POLR1A associated with none, one, or multiple heterochromatin clusters. (**D**) Representative images POLR1A (red) localized to the nucleolus (demarcated by nucleolin (green)) acquired using immunofluorescent widefield microscopy. (**E**) bar plot summarizing ChIP peak signal for POLR1A and POLR2A in and outside of NADs in mouse ESC cells. (**F**) Spearman correlation for peaks in (**E**) showing POLR2A and POLR1A peak correlation. (**G**) Chromosomal distribution of POLR1A ChIP peaks in HMEC cells. (**H**) rDNA coverage profile for POLR1A, POLR2A, H3K9me3, and H3K4me3 Cut&Tag. (**I**) Coverage profile of POLR1A, POLR2A, H3K9me3, and H3K4me3 Cut&Tag at the TSS of all genes. (**J**) Coverage profile of POLR1A, POLR2A, H3K9me3, and H3K4me3 Cut&Tag at POLR1A peaks called with MACS2 in and outside of NADs.

Finally, we generated Cut&Tag to confirm that Pol-I was bound to extranucleolar chromatin in HCT116 cells. To validate our POLR1A antibody in this context, we mapped POLR1A reads to the 45s ribosomal DNA region (rDNA) using a genome prepared with a single rDNA repeat^50^. As expected, POLR1A signal was enriched at rDNA along with H3K9me3 (**Fig 6h**). Within gene bodies, POLR1A peaks were confined primarily to 5’ end of genes near the TSS or the transcriptional end site. Gene promoters, rich in POLR2A and K4me3 signal, were accompanied by modest POLR1A signal, indicating that Pol-I and Pol-II co-occupy transcriptional start sites (**Fig 6i**). We also aggregated Cut&Tag signal in and outside of NADs at POLR1A peaks. Concordant with previous observations that NADs are in a heterochromatic state, NADs were enriched in signal from H3K9me3^44,45^. POLR1A peak signal was roughly proportional in and outside of NADs, while POLR2A and K4me3 signal at POLR1A peaks were only enriched outside of NADs (**Fig 6j**). These results collectively point towards the intriguing likelihood that Pol-I is non-randomly positioned outside nucleoli at gene loci and co-occupies transcriptional start sites with Pol-II.

### Pol-I coregulates non-nucleolar mRNA expression by restricting intronic and intergenic transcription

The complementary functions of Pol-I and Pol-II on *in situ* chromatin domain structure, heterochromatin organization, genome connectivity, and transcription that we observed gave rise to an unexpected hypothesis: extranucleolar Pol-I may co-regulate chromatin domain organization alongside Pol-II to facilitate coherent genome-wide gene transcription. We reasoned that extranucleolar Pol-I may be engaged in spurious transcripton of non-coding DNA proximal to Pol-II transcribed genes. Pol-I transcribes at a an elevated rate compared to Pol-II and shares many of the same subunits^51^. Likewise, prior work has demonstrated that Pol-I can utilize Pol-II or Pol-III transcriptional machinery^38,53,54^. To test this hypothesis, we performed read-mapping and annotation of our transcriptomic data for intronic and intergenic segments proximal to gene bodies. Total intronic counts were then calculated by the difference in reads for the whole gene body compared to the exon reads as described in Lee et al^54^. Partitioning genes by their exon/intron transcriptional change relative to baseline, we observed a marked down-regulation of intron and exon transcription in the ActD treated group (**Fig SI 7a**).

In contrast to ActD treatment, individual POLR1A depletion and POLR2A depletion resulted in a reciprocal transformation of expression: POLR1A degradation mutually upregulated introns and downregulated exons (**Fig SI 7b**), while POLR2A degradation diminished intronic expression but upregulated exons (**Fig SI 7c**). We calculated Log2 transformed coverage profiles for all samples, binned coverage for all gene bodies, and took the average signal of all bins. ActD intronic coverage decreased to nearly zero, followed by a loss of intronic transcription for POLR2A degraded cells. Remarkably, POLR1A depletion results in the robust genome-wide amplification of intronic coverage. For all three conditions, there was little change in exon coverage (**Fig 7a**).

**Figure 7.**
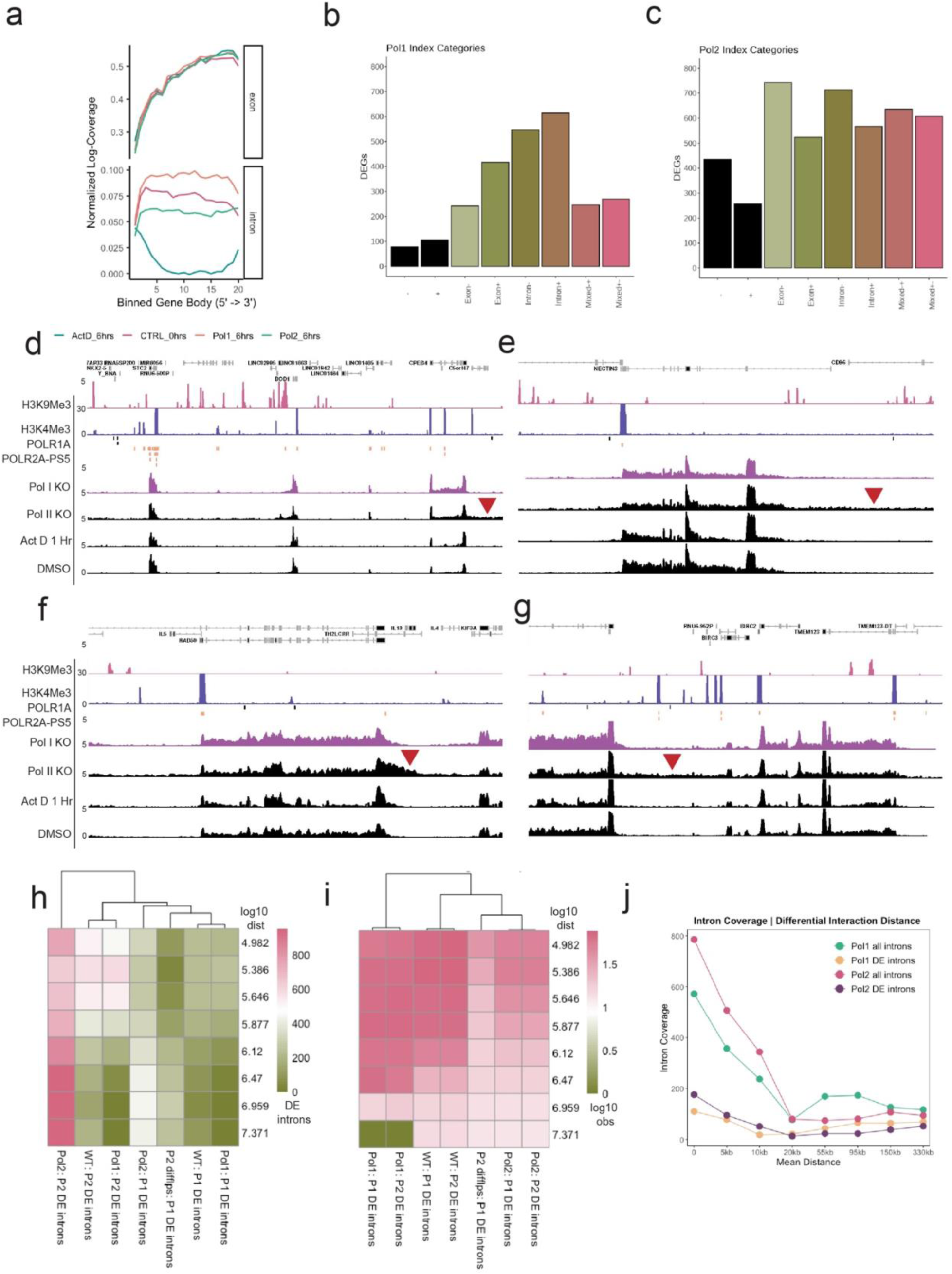
Analysis of intronic and intergenic expression in RNA-seq generated from degron lines (POLR2A-AID2 or POLR1A-AID1) that were treated with auxin for 6 hours or DMSO for 6 hours. (**A**) Intronic signal was binned across all gene bodies and coverage was calculated for each condition. (**B-C**) DEGs for each condition were categorized using Index as (-) downregulated, (+) upregulated, (-exon) downregulated exons, (+exon) upregulated exons, (-intron) downregulated intron, (+intron) upregulated intron, or (+-) mixed for a mixture of the previous categories and summed to generate barplots. (**D-G**) Representative genomic loci where transcriptional readthrough was observed. Red arrows denote readthrough regions. Additional tracks are from publicly available ChIP-seq. Coverage was RPGC normalized and Log2 transformed. (**H**) Loops were binned by distance for each loop set in each DE intron set (denoted Loops:introns) and the number of DE introns were summerd. (**I**) Similar to (**H**) but loop strength at DE intron-loop overlap is summed. (**J**) Differentially interacting regions were discovered using MultiHiCompare; overlap between DI regions and DE introns are plotted as a function of distance.

Intronic RNAs have a faster turnover rate than polyadenylated exonic RNAs and can be used as a proxy to assess nascent RNA synthesis, suggesting that their transformation here was directly related to the loss of each respective polymerase^54,55^. Transcript feature counts were then used to generate analysis of introns and exons for all gene bodies. Consistent with Pol-I and Pol-II co-regulating gene transcription, the loss of POLR2A resulted in a mixed up and down regulation of both introns and exons within the same genes, whereas POLR1A loss resulted in decreased transcription of intronic segments (**Fig. 7b-c**). In the absence of POLR2A, local chromatin organization appears to break down, permitting unrestricted access to introns and exons by other polymerases, generating a mixed transcriptional phenotype.

To characterize polymerase loss at gene loci known to cause pathogenic phenotypes when dysregulated, we generated reads per genome coverage (RPGC) normalized coverage tracks from binary alignment map files for each condition. Coverage scores were transformed to log2-scale for all conditions and offset by 1 to account for 0-values. We analyzed the coverage profiles of several loci that were differentially expressed in both POLR1A degraded and POLR2A degraded cells. In several cases, specific protein coding genes transcribed by Pol-II were downregulated in POLR1A degraded cells, inversely increased in POLR2A degraded cells, and vice-versa. For example, in POLR2A degraded cells, we observed a read-through of exon-intron boundaries and elevated intronic read coverage (**Fig. 7d-g**). Loci in both POLR1A and POLR2A depleted cells revealed extensive 5’ downstream-of-gene intergenic read coverage.

We used publicly available ChIP-seq data to assess the surrounding genomic context of these loci. Notably, while actively transcribing Pol-II is found clustered primarily on exonic regions, Pol-I is sparsely bound at intronic and intergenic regions, consistent with a co-regulatory mechanism in which Pol-I co-organizes and reenforces correct Pol-II transcription of exonal segments for coherent gene expression. We also analyzed PTM marks for euchromatin and heterochromatin, alongside Rad21 and CTCF. Binning up and downregulated genes by log fold change, we quantified average distance from each gene per bin per mark. We found that H3K9me3 heterochromatin distance is positively correlated with gene expression, while H3K4me3 euchromatin mean distance diverges between downregulated and upregulated genes. Unexpectedly, we observed that Rad21 and CTCF mean distance actually diverges for downregulated versus upregulated genes: Rad21 and CTCF are farther away from genes that are downregulated while the more upregulated a gene becomes the closer Rad21 and CTCF are (**SI 7d-e**). Degradation of Rad21 had previously been shown to trigger differential expression when assayed by nascent RNA sequencing^7^.

While large-scale topological contact behavior did not undergo significant reorganization, we previously documented changes to both observed loop strength and size in transcriptionally disrupted cells. To better understand how function is related to contact toplogy, we counted the number of differentially expressed introns in either POLR1A or POLR2A degraded cells that overlapped with loop anchors in both conditions. We also used HiCcompare with standard parameters to find differentially expressed loops in POLR2A degraded cells. We chose to focus on introns for this analysis because intronic expression is a closer approximaton of nascent expression^55^. While DE intron coverage was strongest at short distances in Pol-I loops (between 200 and 300 DE introns per bin), unexpectedly, DE intron coverage over Pol-II loop anchors increased steadily with loop length (700-800 DE introns per bin) (**Fig. 7h**). Observed strength loop strength for each condition was comparable to all loops, irrespective of overlap with DE introns (**Fig. 7i**) (**Fig. 4d**). Finally, we analyzed the relationship between contact behavior and expression by binning the distances between differential interactions found in our data using HiCcompare and counting the overlap with DE introns. We found that DE intron coverage was highest at short distances for differential interactions. DE Intron coverage was enriched compared to coverage for all introns, confirming the relationship between contact behavior and expression for Pol-I and Pol-II (**Fig. 7j**)^56^.

To verify that this exon ‘slippage’ phenotype was reproducible in other data, we analyzed the coverage profiles of the same loci in recently deposited PRO-seq data for Pol-II depleted HCT-116 cells on ENCODE. Consistent with our coverage results, we observed the same readthrough phenotype (**SI 7f-i**). Of these loci, a particularly striking example was the read-through events occurring between RAD50 and IL-13 forward to IL-4 on Chromosome 5 (**Fig. 7f**) (**SI 7f-i**). These genes separately span crucial DNA repair mechanisms (RAD50) and tissue response to inflammatory signals (IL-13 and IL-4)^57–59^. Given the role of IL-13 and IL-4 in inflammatory and atopic disorders, such as asthma, atopic dermatitis, allergic rhinitis, eosinophilic esophagitis, and ulcerative colitis, these results suggested further work is necessary to understand the polymerase-mediated regulatory mechanisms governing chromatin structure between neighboring genes in human disease.

## Discussion and Conclusion

Many questions persist on the interdependent relationship between supranucleosomal 3D chromatin structure and transcription. Conflicting reports from previous studies depend heavily on the mode of investigation. For example, in older chromosome capture studies, degrading Cohesin or CTCF does not disrupt transcription, while depletion of all three polymerases had little effect on largescale features such as TADs and compartments, suggesting that topological features and transcription are independent^4–6^. More recent studies using high resolution chromatin capture modalities and nascent RNA-sequencing have demonstrated that loss of TADs alongside pharmacological inhibition does lead to transcriptional and topological dysregulation, with the caveat that these studies are limited to fine-scale features^7,44^. Largescale features such as TAD boundaries do exert influence over transcriptional output by insulating-enhancer promoter interactions, however, this behavior is largely probabilistic^60^. Similarly, A and B compartments, long thought to insulate transcriptionally active genes from inactive genes, have both been found to harbor actively transcribing genes using RNA-DNA contact mapping^61^. By contrast, super resolution STORM imaging found POL-II distributed along the periphery of chromatin domains throughout the nucleus^3^, alongside euchromatic post-translation modifications on the outside of heterochromatic cores^11^. In super resolution imaging of chromatin, disruption of transcription has a profound impact on 3D chromatin structure^9,10,16^.

To bridge the gap between modes of investigation, we took advantage of an integrative approach described by Li et al in this study; we paired *in situ* structural measurements (PWS nanoscopy, multiplexed SMLM, and ChromSTEM) with gene expression analysis and Hi-C to explore the impact of degrading POLR1A and POLR2A on interphase chromatin domain structure and subsequent function (nano-ChIA). We show that RNA polymerase I is required for coherent mRNA transcription through maintaining the boundaries of genes. Consequently, POLR1A depletion results in dissonant transcriptional signatures where exonic and intronic transcription decouple, characterized by intronic upregulation genome-wide. Additionally, we demonstrate that loss of POLR2A does not lead to exclusive gene downregulation as anticipated; instead POLR2A loss leads to dysregulated expression of exons and introns characterized by frequent readthrough events at gene and exon boundaries.

The functional dysregulation described in this study was accompanied by several organizational changes: we observed a loss of weak distal loops and gain of short-range strong loops in POLR1A degraded cells (**Fig 4b**). Conversely, POLR2A loss precipitated a dramatic increase of weak entropic long-range loops (**Fig 4c**). In both cases, loop anchors overlapped with differentially expressed introns. In the surrounding organizational context, POLR1A depletion triggered heterochromatin core expansion (H3K9me3) while lowering the likelihood that chromatin would be organized into coherent domains (**Fig. 5b,e Fig. 4i-k**). Unexpectedly, ground truth measurement of chromatin organization by chromSTEM tomography revealed that POLR1A loss abruptly triggered chromatin domain degradation, consistent with PWS quantification (**Fig 5j-l**). Here, we also show that Pol-I is spatially situated on the outside of heterochromatic cores throughout the nucleus, not only at sites of rDNA transcription. We confirmed this using publicly available ChIP-seq data and Cut&Tag data, discovering that POLR1A is bound genome-wide.

Our work indicates that much is yet to be understood about how polymerases spatially and functionally cooperate to generate coherent transcripts from coding and non-coding features. The canonical role of Pol-I described in literature has been restricted to rDNA transcription. In comparison, Pol-II is widely understood as solely resonsible for transcription of the majority of the coding and non-coding genome. Some recent studies have challenged this paradigm that each polymerase is spatially and functionally restricted. For instance, Pol-II transcription of non-coding RNAs within the 45s rDNA repeat regulate Pol-I rDNA transcription, indicating that the nucleolus is potentially under a similar co-regulatory polymerase framework^62^. While focused on the co-regulatory role of Pol-II and RNA polymerase III (Pol-III), one study found that degrading Pol-I led to Pol-II led to differentially binding across hundreds of sites^40^. Using RNA-DNA contact mapping, genes actively transcribed by Pol-II were found enriched in contacts to nucleolar organizing hubs^61^. Pol-III consensus binding profiles generated from multiple ChIP datasets predicted widespread Pol-III binding at mRNA promoters^63^. Finally, heterochromatic NADs, rich in cis interchromosomal interactions, overlapped extensively with Lamin Associated Domains (LADs) and appear to be widely distributed across the genome^44,64^.

Together, previous studies along with our findings point towards a reciprocal model: Pol-I may function as a boundary element that, like CTCF, provides 1D sequence specificity to chromatin structure. When no longer constrained by Pol-II, Pol-I freely transcribes through exons and gene body boundaries. The process of unrestricted transcription in turn generates local strong focal enrichment between promoters and enhancers, decompacting chromatin domains and heterochromatin cores in the process. Conversly, without Pol-I providing a counterweight, Pol-II spuriously transcribes through introns, generating weak distal loops (**Fig. 8**). In this model, noncoding elements may function as crucial space-filling scaffolds that form the cores of chromatin domains; these may be maintained by Pol-I to optimize the physical and molecular conditions for Pol-II transcription on coding elements near the surface. In this perspective, read-through behavior accompanied by POLR1A depletion could act as a mechanism for rapid phenotypic switching during stress response by exposing new genomic elements for expression. Supportive prior work suggests that chromatin domain organization acts as a regulator of an integrated stress response for cells to explore their available genomic information by controlling transcriptional heterogeneity and phenotypic plasticity^12,13^. In line with this, future studies focused on identifying the structural principles governing intronic and intergenic transcription may help uncover new patterns in human diseases.

**Figure 8.**
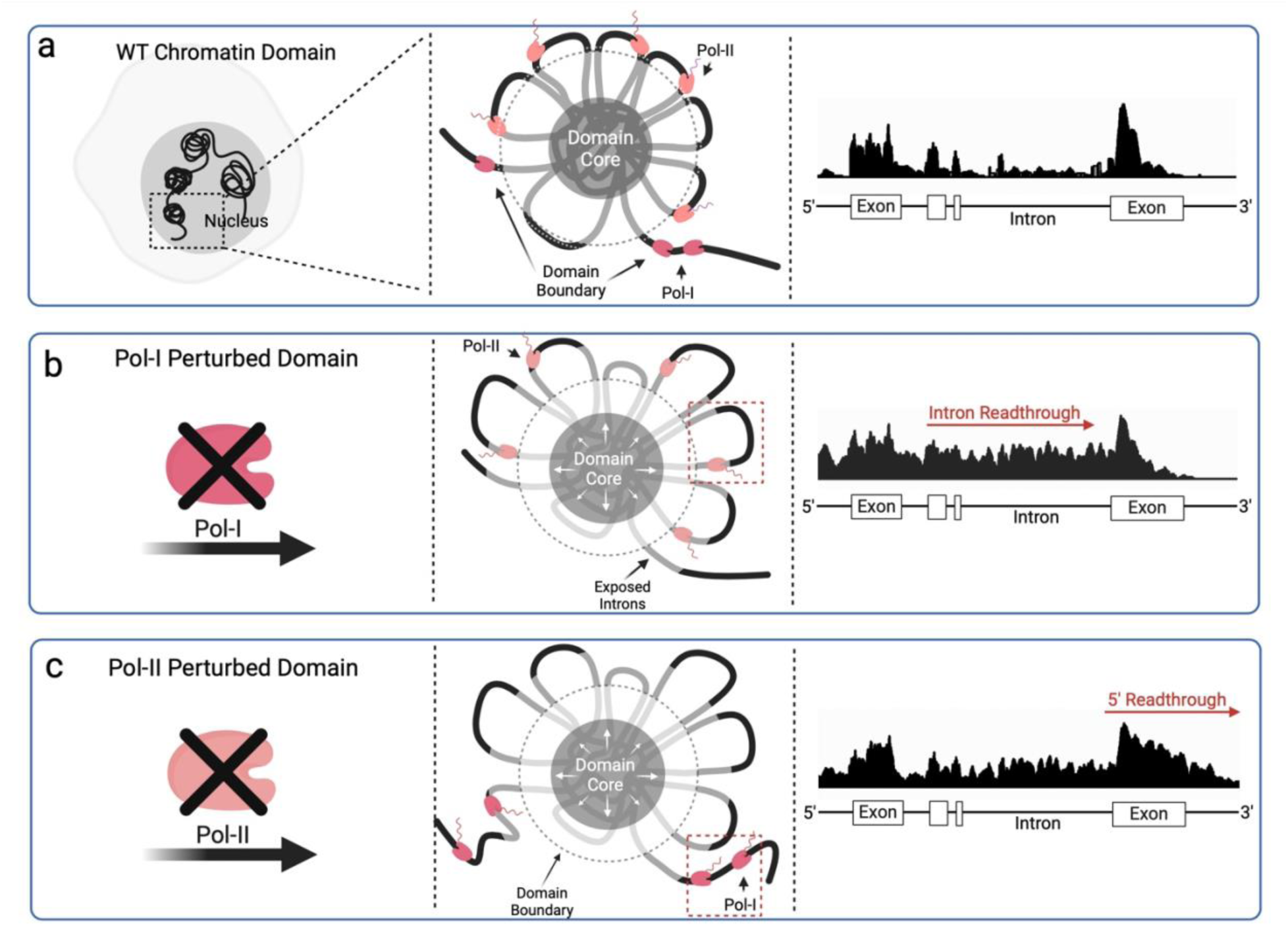
Model for role of Pol-I as a boundary element and functional consequence in loss of Pol-I. (**A**) Wildtype domains where introns are sterically excluded from transcriptional proteins while exons are exposed. Pol-I sits on outside of domain and gene boundaries. (**B**) Chromatin domain dysregulation when Pol-I is lost leads to heterochromatin core expansion and exposure of intronic sequences, resulting in increased Pol-II exon readthrough. (**C**) Chromatin domain dysregulation when Pol-II is lost leads to heterochromatin core expansion and gain of 5’ gene readthrough into intergenic regions.

With this in mind, new questions arise on the genome structure-function relationship 1) How do Pol-I and Pol-II coordinate their positional activity? 2) Do Pol-I and Pol-III act as a short-term failsafes in the absence of Pol-II function or does this represent a method for cells to explore novel transcriptomic space? 3) What is the role of Pol-III in domain organization and co-transcriptional regulation? (4) Do intronic and intergenic sequences demarcate genes with distinct functions (RAD50 and IL-13) within domains to coordinate stress-responses? For example, the read-through of RAD50 (associated with DNA damage response) appears to provide a physical basis to couple damage repair to immune recruitment by the expression of IL-13 (a key cytokine involved in allergic and autoimmune disorders) through the loss of Pol-I. And (5) Pol-I dysregulation is a common feature of multiple cancers; is Pol-I dysregulation mechanistically associated with cancer transcriptional plasticity and heterogeneity? If so, what are the implications for understanding and treating cancers if they can utilize Pol-I mediated transcriptomic sampling of non-coding elements not available to normal cells? We hope that future work utilizing inducible degron systems, paired studies to other chromatin modifying enzymes, and clinical studies will shed light on these evolving questions.

## Limitations of this study

This study focuses on the coregulatory role of RNAPI and RNAPII. These polymerases together account for ∼85% of cellular transcription. It is possible that RNAPIII also contributes to the transcriptional dysregulation phenotype we observed in our data. Studies going forward should utilize the auxin inducible degron system to target RNAPIII, in addition to the other two polymerases. Additionally, although Actinomycin D approximates the phenotype we expect to see when we disrupt all three polymerases, Act D pleiotropy is a poor approximation of polymerase degradation. More complex methods are needed to completely ablate transcription from all three polymerases in a targeted manner and probe the relationship between transcription and chromatin organization. To this end, future studies should generate double degrons targeting pairs of each polymerase.

## Acknowledgements

Philanthropic support was generously received from Rob and Kristin Goldman, the Christina Carinato Charitable Foundation, Mark E. Holliday and Mrs. Ingeborg Schneider, and Mr. David Sachs. This research was supported in part through the computational resources and staff contributions provided for by the Quest high performance computing facility at Northwestern University which is jointly supported by the Office of the Provost, the Office for Research, and Northwestern University Information Technology. This research was supported in part through the computational resources and staff contributions provided by the Genomics Compute Cluster which is jointly supported by the Feinberg School of Medicine, the Center for Genetic Medicine, and Feinberg’s Department of Biochemistry and Molecular Genetics, the Office of the Provost, the Office for Research, and Northwestern Information Technology. The Genomics Compute Cluster is part of Quest, Northwestern University’s high performance computing facility, with the purpose of advancing research in genomics. We appreciate the generous support from the ENCODE Consortium in the generation and dissemination of publicly available datasets. We specifically want to thank the labs of John Lis, Bas van Steensel, Bradley Bernstein, and Richard Myers for the generation and publication of the data utilized within this manuscript. We also want to thank Shelby Blythe, Yue Yang, and Xiaozhong Wang at Northwestern University for their guidance on this project. A special thanks to the lab of Kazuhiro Maeshima for providing us with the POLR1A-AID1 HCT116 degron line.

## Funding

National Institutes of Health grant U54 CA268084

National Institutes of Health grant R01CA228272

National Institutes of Health grant U54 CA261694

NIH Training Grant NIH: T32 GM142604

NIH Training Grant T32AI083216 (LMA)

NSF: EFMA-1830961

The Center for Physical Genomics and Engineering at Northwestern University

## Data and materials availability

All of the data, code used for analysis, and relevant materials have been uploaded to the following locations. Publicly available ChIP-seq, Dam-ID, or Pro-Seq data were obtained from the ENCODE Consortium with the relevant accession numbers are shared in the supplementary information in **Table S1**. Code used for analysis and relevant compiled data to regenerate the figures has been uploaded to GitHub at https://github.com/BackmanLab/Transcription. Raw and processed image data are available upon request. All sequencing data has been deposited at GEO under the following accession number:

## Materials and Methods

### Cell Culture

HCT116-POLR2A-AID2^15^, HCT116-POLR1A-AID^17^, and HCT116 cells were grown in McCoy’s 5A Modified Medium (#16600-082, Thermo Fisher Scientific) supplemented with 10% FBS (#16000-044, Thermo Fisher Scientific) and penicillin-streptomycin (100 μg/ml; #15140-122, Thermo Fisher Scientific). All cells were cultured under recommended conditions at 37°C and 5% CO2. Lines in this study were maintained between passage 5 and 20. Cells were allowed at least 48 hours to re-adhere and recover from trypsin-induced detachment prior to experiments. All experiments and imaging wer performed at a confluence between 40–70%. All cells were tested for mycoplasma contamination (ATCC, #30-1012K) before starting perturbation experiments, and they have given negative results.

### Auxin Treatment and Drug Treatments

HCT116-POLR1A-AID cells or HCT116-POLR2A-AID2 were plated and grown to a confluence between 40–70% prior to treatment. To induce expression of OsTIR1 in HCT116-POLR1A-AID, 1 μg/ml of doxycycline (Fisher Scientific, #10592-13-9) was added to cells 24 hours prior to auxin treatment. HCT116-POLR1A-AID cells were treated with 500 μM Indole-3-acetic acid sodium salt (IAA, Sigma Aldrich, #6505-45-9) to degrade endogenous POLR1A. HCT116-POLR2A-AID2 cells were treated with 1 µM 5-phenyl-indole-3-acetic acid (5-Ph-IAA; MedChemExpress HY-134653) to degrade endogenous POLR2A. HCT116 “WT” cells were treated with either 5 µg/mL Actinomycin D (#0210465810, MP Biomedicals) or % v/v DMSO (#J66650.AE, Thermo Scientific) equal to the highest DMSO concentration in each experiment. Treatment times varied between 1 hour and 8 hours.

### High Throughput Chromatin Conformation Capture

#### Cell culture and sample preparation

Cultured cells were treated with 1 µM 5-PH-IAA (MedChemExpress HY-134653) for 6 hours or with DMSO at an equivalent concentration for 6 hours. At 6 hours, cells were harvested at 1.2 x 10^6^ cells per sample and transferred to a 50 mL Falcon tube, where they were centrifuged for 10 minutes at 450 x g at room temperature. Supernatant was removed and cells were resuspended in 20 mL of cold fresh media. At this point, 540 µL of 37% formaldehyde (#47608, Sigma) was added to bring the final concentration to 1% formaldehyde and fix the cells for 10 minutes. Upon fixation, the reaction was quenched with 10 mL of 3 M Tris (pH 7.5) (#10708976001, Sigma) for 15 minutes. Cells were then centrifuged for 10 minutes at 800 x g and 4 ° C. After resuspension at the desired concentration, individual samples were snap frozen in liquid N2.

#### Hi-C library generation

To wash the nuclei, samples were thawed and resuspended in 50 µL ice cold PB, followed by the addition of 150 µL ice-cold RNase-free water. 50 µL of Buffer C1 (Qiagen Epitect Hi-C Kit: #59971, Qiagen) was added to each sample and mixed. Samples were then centrifuged at 2500 x g and 4 ° C for 5 minutes. The supernatant was aspirated and the nuclear pellet was resuspended in 500 µL of RNase-free water, before being centrifuged again at 2500 x g and 4 ° C for 5 minutes. Library generation and subsequent steps used proprietary reagents from Qiagen’s Epitect Hi-C Kit. Washed nuclei were digested with according to the Epitect Hi-C protocol using a proprietary enzyme cocktail that cut at the GATC motif. Nuclei were end-labeled with biotin followed by ligation for 2 hours at 16 ° C. Ligated chromatin was then de-crosslinked using 20 µL of Proteinase K solution at 56 ° C for 30 minutes and then 80 ° C for 90 minutes. Ligated, decrosslinked DNA was purified using a Qiagen column kit (#59971, Qiagen). and resuspended in 130 µL EB.

#### Library fragmentation

Hi-C library samples were fragmented to a median size of between 400 and 600 bp using a Covaris E220 sonicator with a sample size of 130 µL and the following settings:

|  |  |
| --- | --- |
| Water levels | 12 |
| Peak incident power | 140 W |
| Duty factor | 10% |
| Cycles per burst | 200 |
| Treatment time | 55 seconds |

Samples were purified for fragments between 400 and 500 bp using a Qiagen bead purification size exclusion kit.

#### Hi-C sequencing library generation

Hi-C samples were streptavidin-purified to enrich for properly ligated contact pairs using streptavidin beads and a magnetic bead rack. Beads were first washed in 100 µL of bead wash buffer, resuspended in 50 µL of bead resuspension buffer, and then mixed with 50 µL of Hi-C sample. The mixture was then incubated at room temperature for 15 minutes in a thermal mixer at 1000 RPM. Enriched bead-bound DNA was then end-repaired, phosphorylated, and poly-A tailed using a combined ER/A-tailing solution. The samples were incubated for 15 minutes at 20 ° C followed by incubation at 65 ° C for 15 minutes. Beads were then washed once with 100 µL of bead wash buffer, washed again with 95 µL of adapter ligation buffer, and resuspended in adapter ligation buffer in preparation for ligation of Illumina adapter sequences. Each sample was mixed with 5 µL of one of 6 Illumina adapter sequences (specified in the Qiagen protocol appendix) and 2 µL of ultralow input ligase. These samples were then incubated for 45 minutes. Following adapter ligation, each sample was washed three times before adding 400 µL of library amplification mixture to the beads. Samples were then distributed equally across 8 wells of a 96-well PCR plate and cycled using the following parameters on an Eppendorf Mastercycler x50s thermocycler:

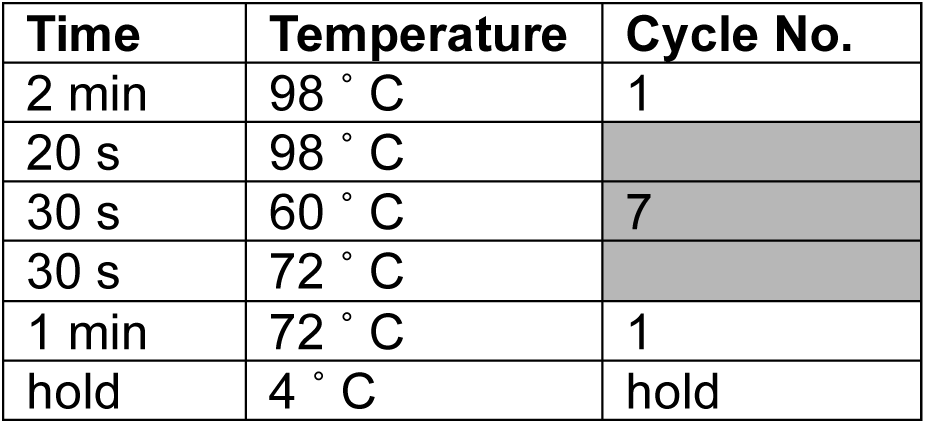

| Time | Temperature | Cycle No. |
| --- | --- | --- |
| 2 min | 98 ° C | 1 |
| 20 s | 98 ° C |  |
| 30 s | 60 ° C | 7 |
| 30 s | 72 ° C |  |
| 1 min | 72 ° C | 1 |
| hold | 4 ° C | hold |

Following sequencing, PCR reactions for each sample were pooled and cleaned using a Qiagen QIAseq library purification kit. Library quality was assessed using a High Sensitivity DNA Assay on an Agilent Bioanalyzer 2100 and a Qubit dsDNA High Sensitivity Assay. Libraries were quantified using a KAPA ROX Low Master Mix qPCR library quantification kit on a QuantStudio 7 Flex instrument. Two samples per lane were sequenced on an Illumina NovaSeq 6000, generating between 400 and 600 million 150 bp paired end reads per sample at Northwestern University’s NUseq Core Facility.

#### Hi-C data processing and analysis

Act D treated samples and 5-ph-IAA treated samples were aligned to human reference genome GRCh38 (Ensembl) using Borrow-Wheeler Aligner version 0.7.17 and processed using the SLURM version of Juicer version 1.6^65^ (https://github.com/aidenlab/juicer) on the Quest HPC provided by Northwestern University. Processing included in the Juicer pipeline included removal of duplicates, exclusion of improperly ligated fragments, and mapping of Hi-C contacts with the GATC motif. Statistics generated for each replicate can be found in the SI. Individual replicates were checked for reproducibility using standard heuristics and combined as a mega map using Juicer’s Mega script to increase sample resolution. TAD identification was generated using Juicer’s Arrowhead (https://github.com/aidenlab/juicer/wiki/Arrowhead). Loops were identified using Juicer’s HICCUPS (https://github.com/aidenlab/juicer/wiki/HiCCUPS). Compartment eigenvector analysis and Pearson correlation analysis were generated using Juicer’s Eigenvector and Pearsons scripts, respectively, or using built in functions in GENOVA. Aggregate TAD Analysis and Aggregate Peak Analysis were generated using GENOVA^66^ (https://github.com/robinweide/GENOVA). Contacts were dumped using Hi-C Straw (https://github.com/aidenlab/straw) and used for downstream analysis. Visualization was also done using GENOVA. Differential interaction analysis and cis-trans interaction analysis was done with the MultiHiCompare package in R^56^ ([CITATION](https://academic.oup.com/bioinformatics/article/35/17/2916/5298730?login=false)). Replicate reproducibility was quantified using the HiCRep package in R^42^ ([CITATION](https://pubmed.ncbi.nlm.nih.gov/28855260/)). All other analyses were custom generated in R. All code used in this publication is available on Github at: https://github.com/BackmanLab/Transcription

### Multi-Color and Single Color SMLM Sample Preparation

#### For H3K9me3 single label imaging

1. Cells were plated on No. 1 borosilicate bottom eight-well Lab-Tek Chambered cover glass (Thermo Scientific, #155411) at seeding density of 12.5k. After 48 hours, the cells underwent fixation for 10 mins at room temperature with a fixation buffer composed of 4% paraformaldehyde (Electron Microscopy Sciences, #15710) in PBS. Samples were then washed in PBS for 5 minutes and then quenched with freshly prepared 0.1% sodium borohydride in PBS for 7 min. Two more wash steps were performed after quenching.
2. Permeabilization was done with blocking buffer composed of (3% bovine serum albumin (BSA) (Fisher, #BP671-10), 0.5% Triton X-100 (Thermo Scientific, #A16046-AE) in PBS) for 1 hour and then samples were immediately incubated with rabbit anti-H3K9me3 (Abcam, ab176916) in blocking buffer for 1-2 hours at room temperature and shaker. Samples were then washed three times with a washing buffer composed of 0.2% BSA, 0.1% Triton X-100 in PBS.
3. Afterwards, samples were incubated with the corresponding goat antibody–dye conjugates, anti-rabbit AF647 (ThermoFisher, A-21245) for 40-60 mins at room temperature on the shaker. After incubation, samples were washed two times in PBS for 5 mins on a shaker and then were ready to be imaged.

#### For H3K9me3 and RNAP polymerase II dual-label imaging

Steps 1,2 and 3 were the same as described above. Samples were then incubated overnight in a modified version of the blocking buffer, which comprises of 10% goat serum (ThermoFisher, 16210064) and 90% prior composition. After overnight blocking, the samples would then go through the same protocol as in step 3 (primary and secondary antibody incubation) but this time with modified blocking buffer (90% original blocking buffer + 10% goat serum) and washing buffer (99% original washing buffer + 1% goat serum) for the second target. The second target primary antibody was a rat anti-RNA Polymerase II (Abcam, ab252855) and the secondary antibody was goat anti-rat AF488 (ThermoFisher, A-150157). As described before, there was an overnight blocking step in between labels at 4°C. After primary and secondary incubation, the samples are washed three times with PBS and then ready to be imaged or stored at 4°C.

#### For H3K9me3 and RNAP polymerase I dual-label imaging

RNAP polymerase I was first labeled as described in the Immunofluorescence Imaging section. Then the samples were incubated overnight in modified version of the blocking buffer. Following overnight blocking, the samples underwent the same procedure as described in step 3 (primary and secondary antibody incubation), but with modified buffers. For the second target (H3K9me3), the blocking buffer was 90% of the original blocking buffer plus 10% goat serum, and the washing buffer consists of 99% of the original washing buffer with 1% goat serum. The primary antibody for the second target was rabbit anti-H3K9me3 (Abcam, ab176916), and the secondary antibody was goat anti-rabbit AF647 (ThermoFisher, A-21245). After incubation with the primary and secondary antibodies, the samples are washed three times with PBS and are then ready for imaging or storage at 4°C.

#### Single Molecule Localization Data Analysis

Acquired data was first processed using the ThunderSTORM ImageJ plugin^67^ to generate the reconstructed images for visualization via the average shifted histogram method, as well as the localization datasets. Each localization dataset was corrected for drift and subsequently filtered such that remaining data had an uncertainty of less than or equal to 40 nm. Localization coordinates (x,y) were then used in a Python point cloud data analysis algorithm which employed the scikit-learn DBSCAN method (min_pts=3, epsilon=50) to cluster the heterochromatic localizations. Cluster size was determined by the area of the Convex Hull fit of the clustered marks and then normalized relative to a circular cluster with radius of 80 nm. POLR2A or POLR1A density was measured by counting the number of POLR2A or POLR1A in dilated contours of the identified cluster periphery (rings that follow the shape of the cluster) and dividing by the ring area. Association was determined by measuring the number of POLR2A or POLR1A that fall within 5 times the area outside of the cluster relative to all POLR2A or POLR1A localizations. The outside cluster condition signifies and POLR2A or POLR1A that is not within any analysis area and thus is not associated with heterochromatic clusters.

### ChromSTEM

#### Electron Microscopy Sample Preparation

Cells were fixed with 2% paraformaldehyde, 2.5% glutaraldehyde (EM-Grade), 2mM CaCl_2_ in 0.1M sodium cacodylate buffer for 30 minutes at room temperature and 30 minutes at 4°C. The cells were then kept in cold temperature, if possible, for further treatments. After fixation, cells were washed 5x for 2 minutes each wash with 0.1M sodium cacodylate buffer and blocked with 10mM glycine, 10mM potassium cyanide, 0.1M sodium cacodylate buffer for 15 minutes.

The cells were stained with 10μM DRAQ5, 0.1% saponin, 0.1M sodium cacodylate buffer for 10 minutes, followed by 3x washes for 5 minutes each with the blocking buffer. After that, the cells were photo-oxidized in 2.5mM 3,3’-Diaminobenzidine (EM-Grade) under a 100X oil objective, 15W Xenon lamp and Cy5 filter for 5 minutes. The cells were washed 5x times for 2 minutes each with 0.1M sodium cacodylate buffer and stained with 2% osmium tetroxide, 1.5% potassium ferrocyanide, 2mM CaCl_2_ in 0.15M sodium cacodylate buffer for 30 minutes. Cells were washed 5x times for 2 minutes each with Millipore water afterwards.

Cells were then dehydrated gradually with 30%, 50%, 70%, 85%, 95%, 2 times 100% ethanol. After that, cells were incubated under room temperature with 100% ethanol, followed by embedding with Durcupan resin with the standard protocol. After 48 hours of resin incubation at 60° C, the resin blocks were collected for ultramicrotomy.

For ultramicrotomy, an ultramicrotome (UC7, Leica) along with a 35° Diatome knife were used to section 120 nm thick resin samples. The sections were collected on a copper slot grid (2 x 0.5mm) with formvar/carbon film. Gold nanoparticles with 10nm diameter were deposited on both surfaces of the grid afterwards as fiducial markers.

#### Image Collection and Tomography Reconstruction

Images were collected with Hitachi HD2300 STEM microscope at 200kV with HAADF imaging mode, at a magnification of 50kX. Two sets of tilt series images were collected by rotating samples from -60° to +60° at a 2° step, with two roughly perpendicular rotation axes.

For post-processing, IMOD was used to align the images. The gold nanoparticles of the collected images were removed with IMOD for another set of data without the influence of extreme values from gold nanoparticles. For each tilt series, Tomopy was used to reconstruct the volume with Penalized maximum likelihood algorithm with weighted linear and quadratic penalties. The two independent reconstructed volumes were then combined in IMOD, with gold nanoparticles as the matching model and repeated on the data without nanoparticles.

After reconstruction, the top and the bottom 0.1% pixel values were capped to remove extreme values. The pixel values are then scaled between 0 to 1 for analysis.

#### Chromatin Domain Identification and Analysis

Chromatin domains were identified and analyzed following the approach previously described^19^.

For identification of domains, a Gaussian filter with radius = 5 pixels was used followed by CLAHE contrast enhancement on the 2D projection of the 3D tomogram using ImageJ. Chromatin domain centers were identified as local maxima with prominence = 1.5 x standard deviation of pixel values.

To quantify domain properties, an 11 x 11 pixels window is applied to each domain and sampled for each domain. Packing scaling analysis is done by measuring the total intensity of the chromatin that radially expands from the center pixel picked and weighted by the intensity of the center pixel. The linear region of the packing scaling behavior is identified by MATLAB ‘ischange’ function. The domain size is measured as the point at which the packing scaling behavior deviates from the linear fits with 5% difference, or when local packing scaling exponent D reaches 3. CVC of a domain is measured with the average value within the domain on a binarized image with Otsu-binarization algorithm after CLAHE contrast enhancement in ImageJ. The packing efficiency is calculated using the same approach described previously, where I is the average chromatin intensity of a domain, R_f_ is the size of the domain, R_min_ = 10nm is the smallest unit of random chromatin polymer chain and D is the packing scaling exponent of the domain packing behavior^19^. All reagents are listed on **Table S2.**

### Total RNA library preparation and sequencing

Stranded total RNA and small RNA sequencing was conducted in the Northwestern University NUSeq Core Facility. Briefly, total RNA examples were checked for quality on Agilent Bioanalyzer 2100 and quantity with Qubit fluorometer. For sequencing of large RNA species, the Illumina Stranded Total RNA Library Preparation Kit was used to prepare sequencing libraries. The Kit procedure was performed without modifications. This procedure includes rRNA depletion, remaining RNA purification and fragmentation, cDNA synthesis, 3’ end adenylation, Illumina adapter ligation, library PCR amplification and validation. For sequencing of small RNAs, the NEXTFLEX Small RNA-Seq Kit v4 from Revvity was used to build sequencing libraries. First, 3’ RNA adapter and then 5’ adapter were ligated to microRNAs and other small RNAs in the samples. After ligation, a reverse transcription step was carried out to generate single-stranded cDNA. A PCR step is then conducted to amplify the cDNA and incorporate Illumina adapter sequences including index sequences. The amplified cDNA constructs were then purified with a bead cleanup step prior to sequencing. Illumina NovaSeq X Plus and HiSeq 4000 NGS Systems were used to generate single 50-base reads.

### RNA-seq data analysis

RNA-seq reads were preprocessed with FastQC v.0.12.0 and aligned with STAR v.2.6.0^68^ using the --quantMode TranscriptomeSAM parameter to human reference genome GRCh38 (Ensembl). Aligned samples were converted to binary alignment map (BAM) files, sorted, and indexed using SAMtools v.1.6^69^. Coverage files were generated as BigWigs for the postive and negative strand using DeepTools v.3.1.1^70^ BamCoverage utility and normalized by either CPM or RPGC. Coverage files were later merged for ease of analysis. Alignment statistics were generated using SAMtools Flagstat utility. Aligned and processed BAMs were then quantified to generate a counts table using the rsem-calculate-expression command from RSEM v.1.3.3^71^ or the htseq-count command from HTSeq v.2.0.2^72^ and Homo_sapiens.GRCh38.111.gtf (Ensembl) genome annotation. Differential gene expression analysis was performed using DESeq2 v.1.44.0 in R. Intron-centric differential expression analysis was performed using the Intron Differences To Exon or INDEX package (https://github.com/Shians/index) and Superintronic package (https://github.com/sa-lee/superintronic) in R^54^. All code used in this publication is available on Github at: https://github.com/BackmanLab/Transcription.

### ChIP-seq data reanalysis

Chromatin Immuno-precipitation data from GSE145874 on GEO was re-analyzed to assess POLR1A and POLR2A binding profiles in and outside of nucleolar regions. FASTQs were downloaded using the fastq-dump command from SRAtoolkit v.3.0.0. FASTQ quality was assessed using FASTQC v.12.0. Reads were trimmed using Trimmomatic v.0.39 with the following parameters: ILLUMINACLIP:TruSeq3-PE-2.fa:2:30:10 LEADING:3 TRAILING:3 SLIDINGWINDOW:4:15 MINLEN:30;. Reads were aligned to a custom version of mm39 that includes a 45 KB rDNA repeat for mapping to ribosomal DNA^50^ using Bowtie2 v.2.4.5 with the following parameters: -q -X 2000. Sequence alignment map (SAM) files were converted to binary alignment (BAM) files, sorted, and indexed using SAMtools v.1.6. Coverage files were generated as BigWigs for the postive and negative strand using DeepTools v.3.1.1 BamCoverage utility and normalized by CPM. Alignment statistics were generated using SAMtools Flagstat utility. BAMs were merged using SAMtools into a single sorted and indexed BAM for each condition. Peaks were called on each BAM using MACS2 v.2.1.0 with the following parameters: -f BAMPE --nomodel -q 1e-2 --keep-dup all --extsize 200. Peaks were sorted and intersected with a 5% overlap using BEDtools v.2.30.0. Union peak files were analyzed using custom scripts in R that can be found on Github at: https://github.com/BackmanLab/Transcription.

### Dual PWS Imaging

Briefly, PWS measures the spectral interference signal resulting from internal light scattering originating from nuclear chromatin. This is related to variations in the refractive index distribution (Σ) (extracted by calculating the standard deviation of the spectral interference at each pixel), characterized by the chromatin packing scaling (*D*). *D* was calculated using maps of Σ, as previously described^19,28,74^. Measurements were normalized by the reflectance of the glass medium interface (i.e., to an independent reference measurement acquired in a region lacking cells on the dish). This allows us to obtain the interference signal directly related to refractive index (RI) fluctuations within the cell. Although it is a diffraction-limited imaging modality, PWS can measure chromatin density variations because the RI is proportional to the local density of macromolecules (e.g., DNA, RNA, proteins). Therefore, the standard deviation of the RI (Σ) is proportional to nanoscale density variations and can be used to characterize packing scaling behavior of chromatin domains with length scale sensitivity around 20 – 200 nm, depending on sample thickness and height. Changes in *D* resulting from each condition are quantified by averaging over nearly 2000 cells, taken across 3 technical replicates. Live-cell PWS measurements obtained using a commercial inverted microscope (Leica, DMIRB) using a Hamamatsu Image-EM charge-coupled device (CCD) camera (C9100-13) coupled to a liquid crystal tunable filter (LCTF, CRi VariSpec) to acquire monochromatic, spectrally resolved images ranging from 500-700 nm at 2-nm intervals as previously described^19,28,74^. Broadband illumination is provided by a broad-spectrum white light LED source (Xcite-120 LED, Excelitas). The system is equipped with a long pass filter (Semrock BLP01-405R-25) and a 63x oil immersion objective (Leica HCX PL APO). Cells were imaged under physiological conditions (37°C and 5% CO_2_) using a stage top incubator (In vivo Scientific; Stage Top Systems). All cells were given 48 hours to re-adhere before treatment (for treated cells) and imaging.

### Dynamic PWS Measurements

Dynamic PWS measurements were obtained as previously described^28^. Briefly, dynamics measurements (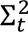, fractional moving mass (*m*_f_), and diffusion) are collected by acquiring multiple backscattered wide-field images at a single wavelength (550 nm) over time (acquisition time), to produce a three-dimensional image cube, where 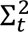 is temporal interference and t is time. Diffusion is extracted by calculating the decay rate of the autocorrelation of the temporal interference. The fractional moving mass is calculated by normalizing the variance of 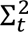 at each pixel. Using the equations and parameters explained in detail in the supplementary information of our recent publication^28^, the fractional moving mass is obtained by using the following equation to normalize 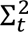 by *ρ*_0_, the density of a typical macromolecular cluster:

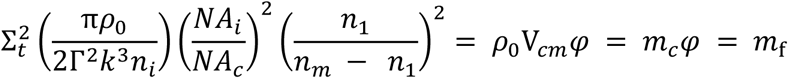

With this normalization, 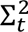 is equivalent to *m*_f_, which measures the mass moving within the sample. This value is calculated from the product of the mass of the typical moving cluster (*m*_*c*_) and the volume fraction of mobile mass (*φ*). *m*_*c*_ is obtained by *m*_*c*_ = V_*cm*_*ρ*_0_, where V_*cm*_ is the volume of the typical moving macromolecular cluster. To calculate this normalization, we approximate *n*_*m*_ = 1.43 as the refractive index (RI) of a nucleosome, *n*_1_ = 1.37 as the RI of a nucleus, *n*_*i*_ = 1.518 as the refractive index of the immersion oil, and *ρ*_0_ = 0.55 g *cm*^-3^ as the dry density of a nucleosome. Additionally, *k* = 1.57E5 cm^-1^ is the scalar wavenumber of the illumination light, and Γ is a Fresnel intensity coefficient for normal incidence. *NA*_*c*_ = 1.49 is the numerical aperture (NA) of collection and *NA*_*i*_ = 0.52 is the NA of illumination. As stated in the aformentioned publication, 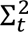 is sensitive to instrument parameters such as the depth of field, substrate refractive index, etc. These dependencies are removed through normalization with the proper pre-factor calculated above for obtaining biological measurements. It should also be noted that backscattered intensity is prone to errors along the transverse direction. Due to these variations, these parameters are more accurate when calculating the expected value over each pixel.

### Coefficient of variation analysis

To assess chromatin compaction through the Coefficient of Variation (CoV) analysis, DAPI-stained cells (see section Fixed-cell immunofluorescence) treated with Auxin (see section “Auxin treatment”) were imaged on a Nikon SoRa Spinning Disk confocal microscope (see section “Confocal imaging”). Following a published workflow, we used ImageJ to create masks of each nucleus. The coefficient of variation of individual nuclei was calculated in MATLAB, with CoV = σ/μ, where σ represents the standard deviation of the intensity values and μ representing the mean value of intensity of the nucleus^74^.

### Immunofluorescence Imaging

HCT116-POLR2A-AID2 and HCT116-POLR1A-AID cells were plated at 10,000 cells per well of an 8-chamber cover glass plate (Cellvis C8-1.5H-N). Following auxin treatment, cells were washed twice with 1x Phosphate Buffered Saline (PBS) (Gibco, #10010031). Cells were fixed with 4% paraformaldehyde (PFA) (Electron Microscopy Sciences, #15710) for 10 minutes at room temperature, followed by washing with PBS 3 times for 5 minutes each. Cells were permeabilized for 15 minutes using 0.2% TritonX-100 (10%) (Sigma-Aldrich, #93443) in 1x PBS, followed by another wash with 1x PBS for 3 times for 5 minutes each. Cells were blocked for 1 hour using 3% BSA (Sigma-Aldrich, #A7906) in PBST (Tween-20 in 1x PBS) (Sigma-Aldrich, #P9416) at room temperature. The following primary antibodies were added overnight at 4°C: Rat POLR2A PS2 antibody (ABCam, AB252855) or Mouse monoclonal RPA194 (Santa Cruz, sc-48385). Cells were washed with 1x PBS 3 times for 5 minutes each. Either of the following secondary antibodies were added for 1 hour at room temperature: Goat Anti-Rat IgG Alexa Fluor 647 (Abcam, ab150167, dilution 1:1000) or Donkey Anti-Mouse IgG H+L Alexa Fluor 647 (Thermo Fisher Scientific, A31572, dilution 1:500). Cells were washed with 1x PBS 3 times for 5 minutes each. Finally, cells were stained with DAPI (Thermo Fisher Scientific, #62248, diluted to 0.5 μg/mL in 1x PBS) for 10 minutes at room temperature. Prior to imaging, cells were washed with 1x PBS twice for 5 minutes each. The cells were imaged using the Nikon Eclipse Ti2 inverted microscope equipped with a Prime 95B Scientific CMOS camera. Images were collected using a 100x/ 1.42 NA oil-immersion objective mounted with a 2.8x magnifier. mClover was excited at 488 nm laser, Alexa Fluor 647 was excited with at 633 nm, and DAPI was excited at 405 nm. Imaging data were acquired by Nikon acquisition software.

*Immunofluorescence imaging for* ***Figure 6d*** *and SI Figure 3c were acquired using the following protocol:* HCT116 POLR1A-AID cells were plated on cover glasses. For **SI Figure 3c**, when cells were 50% confluent, they were treated with doxycycline (1 µg/mL) for 24 hours and then with 500 µm indole-3-acetic acid (IAA) in the presence of doxycycline for 5 hours before being fixed. Cells were chemically fixed with 2% paraformaldehyde (PFA) for 15 minutes, washed with 1x phosphate buffered saline (PBS), then permeabilized with 0.5% triton for 10 minutes, then washed again before overnight primary antibody incubation. The primary antibodies used include RPA194 mouse (Santa Cruz, sc-48385) at 1:200, C23 (nucleolin) rabbit (Santa Cruz, sc-13057) at 1:500, UBF human (provided by Edward Chen at the University of Florida) at 1:600. Cells were washed after primary antibody, then incubated in secondary antibodies: anti-mouse 594 (Invitrogen, A11032) at 1:200, anti-rabbit 488 (Invitrogen, A11008) at 1:200, and anti-human 647 (Invitrogen, A21445) at 1:100. Cells were then washed and mounted with VECTASHIELD antifade mounting medium with DAPI (H-1200). Cells were imaged using Nikon Eclipse Ti2 widefield fluorescence microscope.

### EU-labeled nascent RNA staining and imaging

For EU-labeled nascent RNA staining and imaging in vivo, we used the Click-iT RNA Alexa Fluor 647 Imaging Kit (Thermo Fisher Scientific, C10330) according to the standard immunofluorescence protocol. Cells were plated at 50,000 per well in a 12-well optical plate (Cellvis, P12-1.5H-N) for widefield fluorescence imaging or at 10,000 cells per chamber in an 8-chamber cover glass plate (Cellvis C8-1.5H-N). 48 hours after plating, partially confluent dishes were treated as described above to degrade POLR1A, POLR2A, or all three polymerases (5 µg/mL Actinomycin D). 1 hour prior to fixation, 1 mM EU was added to each well. Cells were washed with PBS and fixed with 4% paraformaldehyde for 10 minutes followed by three more washes in 3% bovine serum albumin (BSA) PBS and permeabilization using 0.5% Triton X-100 in PBS for 20 min at room temperature. Each well was then washed twice with 3% BSA in PBS and incubated with Click-iT reaction mix (Invitrogen) for 30 min. After the Click-iT reaction, cells were washed twice with 3% BSA in PBS. For widefield epifluorescence, all conditions were imaged using the Nikon Eclipse Ti2 inverted microscope equipped with a Prime 95B Scientific CMOS camera quantified with Fiji v.2.14.0. For EU-STORM, all conditions were imaged using the SMLM parameters described previously in this method.

### Protein detection

HCT116-POLR2A-AID2 cells were lysed using Radio Immuno Precipitation Assay (RIPA) buffer (Sigma-Aldrich, #R0278) with protease inhibitor added (Sigma-Aldrich, #P8340). Cell lysates were quantified with a standard Bradford assay using the Protein Assay Dye Concentrate (BioRad, #500–0006) and BSA as a control. Heat denatured protein samples were resolved on a 4–12% bis–tris gradient gel, transferred to a PVDF membrane using the Life Technologies Invitrogen iBlot Dry Transfer System (Thermo Fisher Scientific, IB1001) (20 V for 7 min), and blocked in 5% nonfat dried milk (BioRad, #120–6404) in 1 × TBST. Whole-cell lysates were blotted against the following primary antibodies: anti-POLR2A (Abcam, ab5408, 1:200 dilution). The following secondary antibodies were used: anti-rabbit IgG HRP (Promega, #W4018). Blots were incubated with the primary antibody overnight at 4 °C, followed by incubation with the secondary antibodies for 1 h at room temperature. To develop blots for protein detection, chemiluminescent substrates were used (Thermo Fischer Scientific, #32106). To quantify the western blot bands, we used the iBright Analysis Software from Thermo Fisher Scientific to define bands as regions of interest. By measuring the mean grey intensity values, the final relative quantification values were calculated as the ratio of each protein band relative to the lane’s loading control for all three replicates.

### Flow Cytometry

#### Data and image analysis

Flow cytometry analysis for HCT116-POLR2A-AID2 and HCT116-POLR1A-AID cells to determine proper auxin treatment concentration was performed on a BD LSRFortessa Cell Analyzer FACSymphony S6 SORP system, located at the Robert H. Lurie Comprehensive Cancer Center Flow Cytometry Core Facility at Northwestern University in Evanston, IL. For all FACS analysis, the same protocol was used. After degrading endogenous POLR1A or POLR2A, cells were harvested and analyzed. Briefly, cells were washed with DPBS (Gibco, #14190–144), trypsinized (Gibco, #25200–056), neutralized with media, and then centrifuged at 500 × g for 5 min. Cells were then resuspended in cold FACS buffer (DPBS with 1% of BSA and 2mM EDTA added) at 4°C and immediately analyzed for GFP signal. All flow cytometry data were analyzed using FlowJo software v. 10.6.1.

#### CUT&Tag

CUT&Tag was performed as previously described^75^ using the CUT&Tag-IT Assay Kit (Active Motif, Carlsbad, CA, USA) with modifications recommended in the manufacturer’s protocol. 100,000-200,000 HCT116 cells were harvested at room temperature per replicate. Three biological replicates were prepared on different days for each condition. Antibodies used for Cut&Tag include Mouse monoclonal anti-RPA194 (Santa Cruz, sc-48385), Rabbit polyclonal Histone H3K4me3 antibody (Active Motif, cat # 39916), Rabbit polyclonal Histone H3K9me3 antibody (Active Motif, cat # 39065), Rabbit polyclonal Histone H3K27ac antibody (Active Motif, cat # 39034), and Abflex Mouse RNA Polymerase II antibody (Active Motif, cat # 91152). Samples were sequenced on a single lane for each biological replicate on an Illumina NovaSeq X Plus using 150 BP paired end sequencing at Northwestern University’s NUseq Core Facility.

#### CUT&Tag analysis

Analysis was performed following the recommendations of^76^ (Zheng Y et al (2020). Protocol.io) with some modifications. Reads were trimmed with Trim Galore v.0.6.10 and mapped to the human reference genome GRCh38 (Ensembl) using Bowtie2 v.2.5.4^77^ with the following parameters: --local --very-sensitive --no-mixed --no-discordant -- phred33 -I 10 -X 700. Duplicate reads were marked and removed using Picard v.2.21.4^78^. To account for variation in sequencing depth, once duplicates were removed, replicates were downsampled to the lowest read depth for a given antibody, either 1 million reads, 2 million reads, or 10 million reads. Aligned and dedupped samples were converted to binary alignment map (BAM) files, sorted, and indexed using SAMtools v.1.6. BAM files were converted using Samtools and BED files were converted using Bedtools v.2.30.0^79^. BigWig coverage was generated using Deeptools v.3.1.1 bamCoverage and either CPM or RPGC normalization. BAMs from downsampled replicates were merged for final consensus peak calling. Peaks were called on BAMs with MACS2 v. 2.2.9.1^80^ using the -q 0.1 --keep-dup all parameters and on BEDs with SEACR v.1.3^81^ (^74^) using the 0.01 non relaxed parameters. Matrices of peak coverage profiles were generated using Deeptools v.3.1.1 computeMatrix and heatmaps were plotted using plotHeatmap. Visualizations of coverage files were done IGV^82^.

## References

1. Maeshima, K., Ide, S. & Babokhov, M. Dynamic chromatin organization without the 30-nm fiber. Current Opinion in Cell Biology 58, 95–104 (2019).

2. Dynamic Organization of Chromatin Domains Revealed by Super-Resolution Live-Cell Imaging. vol. 67 (United States, 2017).

3. Li, Y. et al. Nanoscale chromatin imaging and analysis platform bridges 4D chromatin organization with molecular function. Sci Adv 7, eabe4310 (2021).

4. Jiang, Y. et al. Genome-wide analyses of chromatin interactions after the loss of Pol I, Pol II, and Pol III. Genome Biol 21, 158 (2020).

5. Rao, S. S. P. et al. Cohesin Loss Eliminates All Loop Domains. Cell 171, 305–320.e24 (2017).

6. Nora, E. P. et al. Targeted Degradation of CTCF Decouples Local Insulation of Chromosome Domains from Genomic Compartmentalization. Cell 169, 930–944.e22 (2017).

7. Hsieh, T.-H. S. et al. Enhancer-promoter interactions and transcription are largely maintained upon acute loss of CTCF, cohesin, WAPL or YY1. Nat Genet 54, 1919–1932 (2022).

8. Zhang, S. et al. RNA polymerase II is required for spatial chromatin reorganization following exit from mitosis. Sci Adv 7, eabg8205 (2021).

9. Neguembor, M. V. et al. Transcription-mediated supercoiling regulates genome folding and loop formation. Mol Cell 81, 3065–3081.e12 (2021).

10. Kant, A. et al. Active transcription and epigenetic reactions synergistically regulate meso-scale genomic organization. Nat Commun 15, 4338 (2024).

11. Miron, E. et al. Chromatin arranges in chains of mesoscale domains with nanoscale functional topography independent of cohesin. Sci Adv 6, eaba8811 (2020).

12. Virk, R. K. A. et al. Disordered chromatin packing regulates phenotypic plasticity. Sci Adv 6, eaax6232 (2020).

13. Almassalha, L. M. et al. The Global Relationship between Chromatin Physical Topology, Fractal Structure, and Gene Expression. Sci Rep 7, 41061 (2017).

14. Almassalha, L. M. et al. The Greater Genomic Landscape: The Heterogeneous Evolution of Cancer. Cancer Res 76, 5605–5609 (2016).

15. Yesbolatova, A. et al. The auxin-inducible degron 2 technology provides sharp degradation control in yeast, mammalian cells, and mice. Nat Commun 11, 5701 (2020).

16. Nagashima, R. et al. Single nucleosome imaging reveals loose genome chromatin networks via active RNA polymerase II. J Cell Biol 218, 1511–1530 (2019).

17. Ide, S., Imai, R., Ochi, H. & Maeshima, K. Transcriptional suppression of ribosomal DNA with phase separation. Sci Adv 6, eabb5953 (2020).

18. Natsume, T., Kiyomitsu, T., Saga, Y. & Kanemaki, M. T. Rapid Protein Depletion in Human Cells by Auxin-Inducible Degron Tagging with Short Homology Donors. Cell Rep 15, 210–218 (2016).

19. Li, Y. et al. Analysis of three-dimensional chromatin packing domains by chromatin scanning transmission electron microscopy (ChromSTEM). Sci Rep 12, 12198 (2022).

20. Ou, H. D. et al. ChromEMT: Visualizing 3D chromatin structure and compaction in interphase and mitotic cells. Science 357, (2017).

21. Nozaki, T. et al. Condensed but liquid-like domain organization of active chromatin regions in living human cells. Sci Adv 9, eadf1488 (2023).

22. Sanders, J. T. et al. Loops, topologically associating domains, compartments, and territories are elastic and robust to dramatic nuclear volume swelling. Sci Rep 12, 4721 (2022).

23. Li, W. S. et al. Chromatin packing domains persist after RAD21 depletion in 3D. Preprint at 10.1101/2024.03.02.582972 (2024).

24. Snyers, L., Laffer, S., Löhnert, R., Weipoltshammer, K. & Schöfer, C. CX-5461 causes nucleolar compaction, alteration of peri- and intranucleolar chromatin arrangement, an increase in both heterochromatin and DNA damage response. Sci Rep 12, 13972 (2022).

25. Perry, R. P. & Kelley, D. E. Inhibition of RNA synthesis by actinomycin D: characteristic dose-response of different RNA species. J Cell Physiol 76, 127–139 (1970).

26. Bensaude, O. Inhibiting eukaryotic transcription: Which compound to choose? How to evaluate its activity? Transcription 2, 103–108 (2011).

27. Almassalha, L. M. et al. Label-free imaging of the native, living cellular nanoarchitecture using partial-wave spectroscopic microscopy. Proc Natl Acad Sci U S A 113, E6372–E6381 (2016).

28. Gladstein, S. et al. Multimodal interference-based imaging of nanoscale structure and macromolecular motion uncovers UV induced cellular paroxysm. Nat Commun 10, 1652 (2019).

29. Pujadas Liwag, E. M. et al. Nuclear blebs are associated with destabilized chromatin packing domains. Preprint at 10.1101/2024.03.28.587095 (2024).

30. Xu, J., Ma, H. & Liu, Y. Stochastic Optical Reconstruction Microscopy (STORM). Curr Protoc Cytom 81, 12.46.1–12.46.27 (2017).

31. Lelek, M. et al. Single-molecule localization microscopy. Nat Rev Methods Primers 1, 39 (2021).

32. Hollstein, U. Actinomycin. Chemistry and mechanism of action. Chem. Rev. 74, 625–652 (1974).

33. Sobell, H. M. Actinomycin and DNA transcription. Proc Natl Acad Sci U S A 82, 5328–5331 (1985).

34. Lu, D.-F. et al. Actinomycin D inhibits cell proliferations and promotes apoptosis in osteosarcoma cells. Int J Clin Exp Med 8, 1904–1911 (2015).

35. Trask, D. K. & Muller, M. T. Stabilization of type I topoisomerase-DNA covalent complexes by actinomycin D. Proc Natl Acad Sci U S A 85, 1417–1421 (1988).

36. Koba, M. & Konopa, J. [Actinomycin D and its mechanisms of action]. Postepy Hig Med Dosw (Online) 59, 290–298 (2005).

37. Espinoza, J. A. et al. Chromatin damage generated by DNA intercalators leads to degradation of RNA Polymerase II. Nucleic Acids Res 52, 4151–4166 (2024).

38. Watt, K. E., Macintosh, J., Bernard, G. & Trainor, P. A. RNA Polymerases I and III in development and disease. Semin Cell Dev Biol 136, 49–63 (2023).

39. Khatter, H., Vorländer, M. K. & Müller, C. W. RNA polymerase I and III: similar yet unique. Curr Opin Struct Biol 47, 88–94 (2017).

40. Jiang, Y. et al. Cross-regulome profiling of RNA polymerases highlights the regulatory role of polymerase III on mRNA transcription by maintaining local chromatin architecture. Genome Biol 23, 246 (2022).

41. Hyle, J. et al. Auxin-inducible degron 2 system deciphers functions of CTCF domains in transcriptional regulation. Genome Biol 24, 14 (2023).

42. Castells-Garcia, A. et al. Super resolution microscopy reveals how elongating RNA polymerase II and nascent RNA interact with nucleosome clutches. Nucleic Acids Res 50, 175–190 (2022).

43. Yang, T. et al. HiCRep: assessing the reproducibility of Hi-C data using a stratum-adjusted correlation coefficient. Genome Res 27, 1939–1949 (2017).

44. Hsieh, T.-H. S. et al. Resolving the 3D Landscape of Transcription-Linked Mammalian Chromatin Folding. Mol Cell 78, 539–553.e8 (2020).

45. Bersaglieri, C. et al. Genome-wide maps of nucleolus interactions reveal distinct layers of repressive chromatin domains. Nat Commun 13, 1483 (2022).

46. Yu, S. & Lemos, B. The long-range interaction map of ribosomal DNA arrays. PLoS Genet 14, e1007258 (2018).

47. Heinz, S. et al. Transcription Elongation Can Affect Genome 3D Structure. Cell 174, 1522–1536.e22 (2018).

48. Quinodoz, S. A. et al. Higher-Order Inter-chromosomal Hubs Shape 3D Genome Organization in the Nucleus. Cell 174, 744–757.e24 (2018).

49. Peng, T. et al. Mapping nucleolus-associated chromatin interactions using nucleolus Hi-C reveals pattern of heterochromatin interactions. Nat Commun 14, 350 (2023).

50. Maiser, A. et al. Super-resolution in situ analysis of active ribosomal DNA chromatin organization in the nucleolus. Sci Rep 10, 7462 (2020).

51. George, S. S., Pimkin, M. & Paralkar, V. R. Construction and validation of customized genomes for human and mouse ribosomal DNA mapping. J Biol Chem 299, 104766 (2023).

52. Jacobs, R. Q. & Schneider, D. A. Transcription elongation mechanisms of RNA polymerases I, II, and III and their therapeutic implications. J Biol Chem 300, 105737 (2024).

53. Misiaszek, A. D. et al. Cryo-EM structures of human RNA polymerase I. Nat Struct Mol Biol 28, 997–1008 (2021).

54. Russell, J. & Zomerdijk, J. C. B. M. The RNA polymerase I transcription machinery. Biochem Soc Symp 203–216 (2006) doi:10.1042/bss0730203.

55. Lee, S., et al. Covering all your bases: incorporating intron signal from RNA-seq data. NAR Genom Bioinform 2, lqaa073 (2020).

56. Gaidatzis, D., Burger, L., Florescu, M. & Stadler, M. B. Analysis of intronic and exonic reads in RNA-seq data characterizes transcriptional and post-transcriptional regulation. Nat Biotechnol 33, 722–729 (2015).

57. Stansfield, J. C., Cresswell, K. G., Vladimirov, V. I. & Dozmorov, M. G. HiCcompare: an R-package for joint normalization and comparison of HI-C datasets. BMC Bioinformatics 19, 279 (2018).

58. Doran, E. et al. Interleukin-13 in Asthma and Other Eosinophilic Disorders. Front Med (Lausanne) 4, 139 (2017).

59. Chang, X. et al. A genome-wide association meta-analysis identifies new eosinophilic esophagitis loci. J Allergy Clin Immunol 149, 988–998 (2022).

60. Bian, L., Meng, Y., Zhang, M. & Li, D. MRE11-RAD50-NBS1 complex alterations and DNA damage response: implications for cancer treatment. Mol Cancer 18, 169 (2019).

61. Zuin, J. et al. Nonlinear control of transcription through enhancer-promoter interactions. Nature 604, 571–577 (2022).

62. Goronzy, I. N. et al. Simultaneous mapping of 3D structure and nascent RNAs argues against nuclear compartments that preclude transcription. Cell Reports 41, 111730 (2022).

63. Abraham, K. J. et al. Nucleolar RNA polymerase II drives ribosome biogenesis. Nature 585, 298–302 (2020).

64. Rajendra, K. C. et al. Evidence of RNA polymerase III recruitment and transcription at protein-coding gene promoters. Preprint at 10.1101/2024.06.08.598009 (2024).

65. Bersaglieri, C. & Santoro, R. Genome Organization in and around the Nucleolus. Cells 8, 579 (2019).

66. Durand, N. C. et al. Juicer Provides a One-Click System for Analyzing Loop-Resolution Hi-C Experiments. Cell Syst 3, 95–98 (2016).

67. van der Weide, R. H., et al. Hi-C analyses with GENOVA: a case study with cohesin variants. NAR Genom Bioinform 3, lqab040 (2021).

68. Ovesný, M., Křížek, P., Borkovec, J., Svindrych, Z. & Hagen, G. M. ThunderSTORM: a comprehensive ImageJ plug-in for PALM and STORM data analysis and super-resolution imaging. Bioinformatics 30, 2389–2390 (2014).

69. Dobin, A. et al. STAR: ultrafast universal RNA-seq aligner. Bioinformatics 29, 15–21 (2013).

70. Danecek, P. et al. Twelve years of SAMtools and BCFtools. Gigascience 10, giab008 (2021).

71. Ramírez, F., Dündar, F., Diehl, S., Grüning, B. A. & Manke, T. deepTools: a flexible platform for exploring deep-sequencing data. Nucleic Acids Res 42, W187–191 (2014).

72. Li, B. & Dewey, C. N. RSEM: accurate transcript quantification from RNA-Seq data with or without a reference genome. BMC Bioinformatics 12, 323 (2011).

73. Anders, S., Pyl, P. T. & Huber, W. HTSeq--a Python framework to work with high-throughput sequencing data. Bioinformatics 31, 166–169 (2015).

74. Eid, A. et al. Characterizing chromatin packing scaling in whole nuclei using interferometric microscopy. Opt Lett 45, 4810–4813 (2020).

75. Martin, L. et al. A protocol to quantify chromatin compaction with confocal and super-resolution microscopy in cultured cells. STAR Protoc 2, 100865 (2021).

76. Kaya-Okur, H. S. et al. CUT&Tag for efficient epigenomic profiling of small samples and single cells. Nat Commun 10, 1930 (2019).

77. Zheng, Y., Ahmad, K. & Henikoff, S. CUT&Tag Data Processing and Analysis Tutorial. (2020).

78. Langmead, B. & Salzberg, S. L. Fast gapped-read alignment with Bowtie 2. Nature Methods 9, 357–359 (2012).

79. Broad Institute. Picard Toolkit. (2019).

80. Quinlan, A. R. & Hall, I. M. BEDTools: a flexible suite of utilities for comparing genomic features. Bioinformatics 26, 841–842 (2010).

81. Zhang, Y. et al. Model-based Analysis of ChIP-Seq (MACS). Genome Biology 9, R137 (2008).

82. Meers, M. P., Tenenbaum, D. & Henikoff, S. Peak calling by Sparse Enrichment Analysis for CUT&RUN chromatin profiling. Epigenetics Chromatin 12, 42 (2019).

83. Robinson, J. T. et al. Integrative genomics viewer. Nat Biotechnol 29, 24–26 (2011).

