## Supplemental Data for "mRNA expression is co-regulated by non-nucleolar RNA polymerase I"

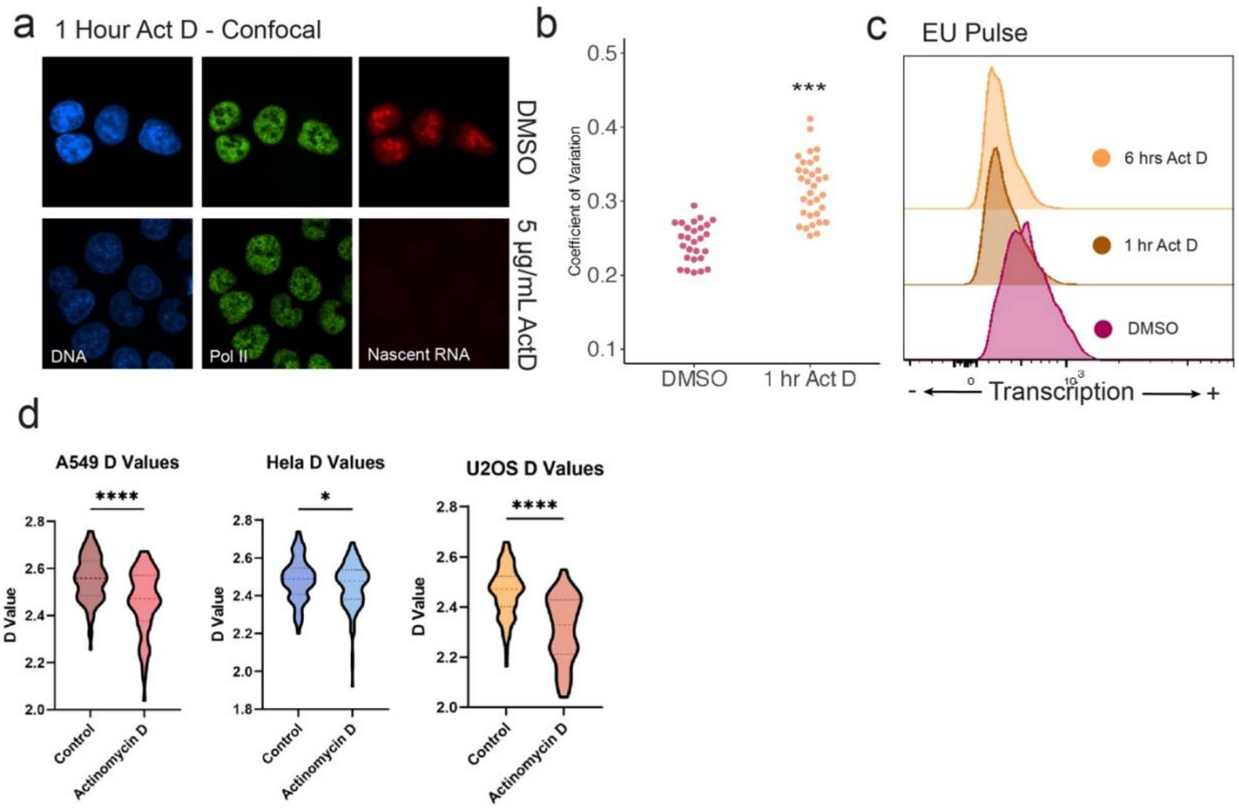


**SI Figure 1.** Treatment with Actinomycin D treated (5 µg/mL) for 1 hour leads to chromatin compaction and abrogation of transcription. (**A**) widefield fluorescent microscopy of ActD treated cells or DMSO control cells: DAPI stained DNA (blue), labelled POLR2A (green), EU-labelled nascent RNA (red). (**B**) Coeefficient of variation analysis for DAPI stained DNA before and after treatment. (**C**) Flow cytometry analysis of EU-labelled nascent RNA before treatment and at 1 or 6 hours of treatment. (**D**) Nuclear scaling analysis of A549, HeLa, and U2OS cells following treatment.

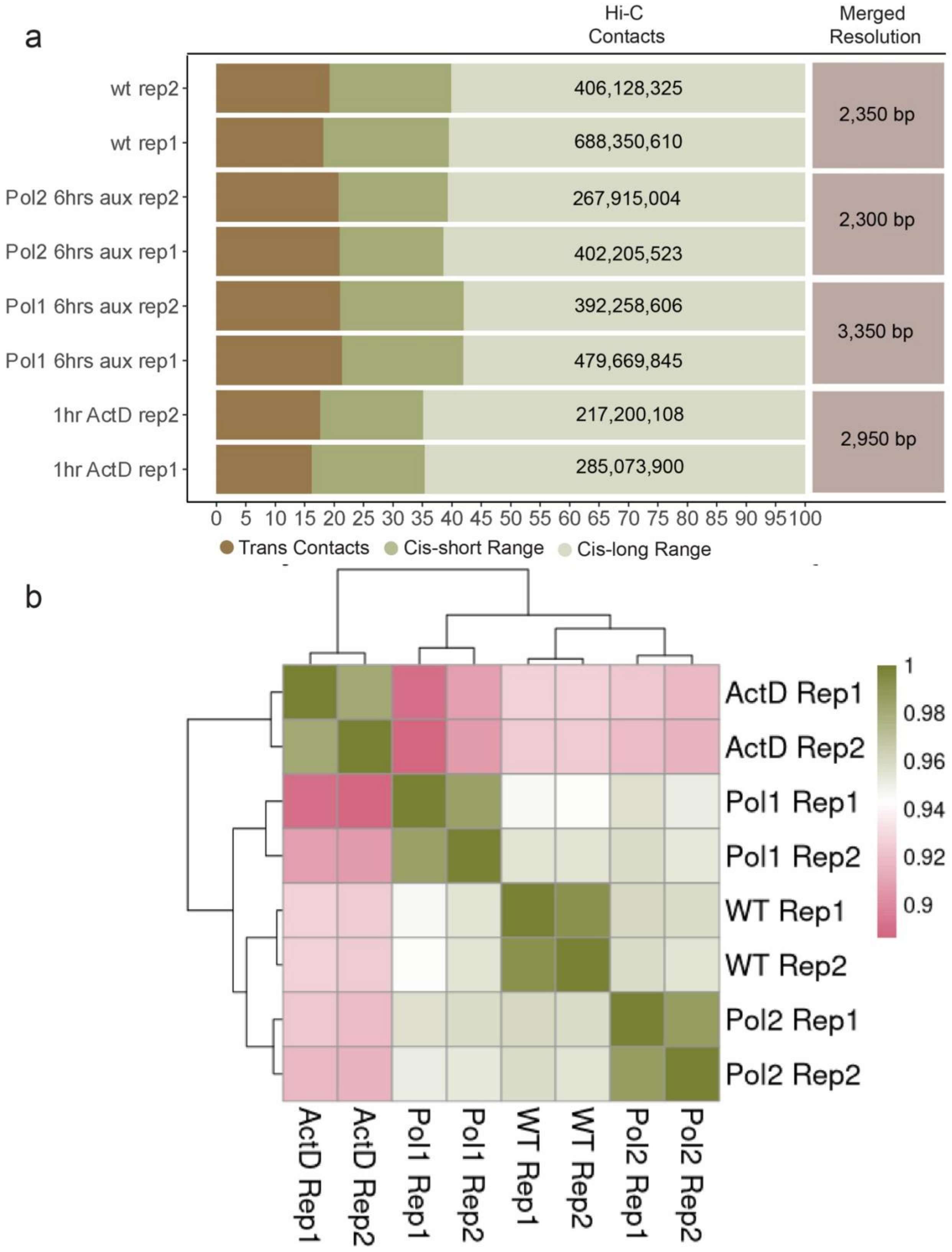
a
Hi-C Contacts
Merged Resolution
wt rep2
wt rep1
Pol2 6hrs aux rep2
Pol2 6hrs aux rep1
Pol1 6hrs aux rep2
Pol1 6hrs aux rep1
1hr ActD rep2
1hr ActD rep1
406,128,325
688,350,610
267,915,004
402,205,523
392,258,606
479,669,845
217,200,108
285,073,900
2,350 bp
2,300 bp
3,350 bp
2,950 bp
0 5 10 15 20 25 30 35 40 45 50 55 60 65 70 75 80 85 90 95 100
Trans Contacts
Cis-short Range
Cis-long Range
b
ActD Rep1
ActD Rep2
Pol1 Rep1
Pol1 Rep2
WT Rep1
WT Rep2
Pol2 Rep1
Pol2 Rep2
1
0.98
0.96
0.94
0.92
0.9

**SI Figure 2.** Two replicates of Hi-C were generated from degron lines (POLR2A-AID2 or POLR1A-AID1) that were treated with auxin for 6 hours, DMSO for 6 hours, or Actinomycin D treated (5 µg/mL) for 1 hour. (**A**) Total number of contacts, trans contacts, and cis contacts for each replicate; resolution of merged replicates. (**B**) The stratum adjusted correlation coefficient (SCC) for all replicates.

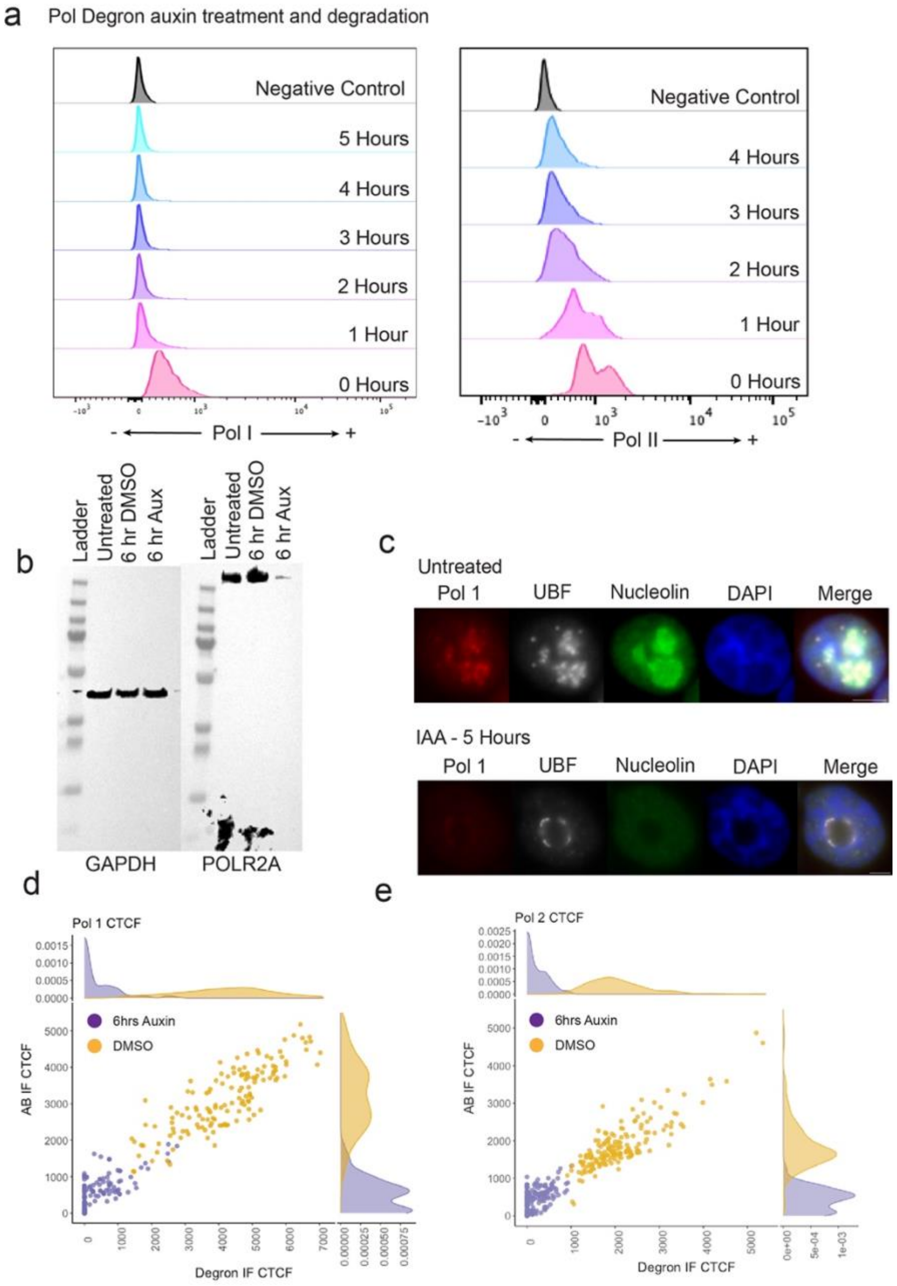

**SI Figure 3.** Validation of HCT116 POLR1A-AID1 and HCT116 POLR2A-AID2 lines. (**A**) Flow cytometry analysis of both lines following treatment with auxin. (**B**) Western blot of POLR2A-AID2 line after 6 hours of degradation. (**C**) POLR1A-AID1 cells were treated with auxin for 5 hours, stained for UBF (white), nucleolin (green), and DNA (blue), and imaged using widefield fluorescent microscopy. (**D-E**) Verification of POLR1A and POLR2A depletion by quantification of degron fluoresence and immunofluorescent staining showing over 90% of the population is without detectable protein. Axes representing corrected-total-fluorescence in relative units.

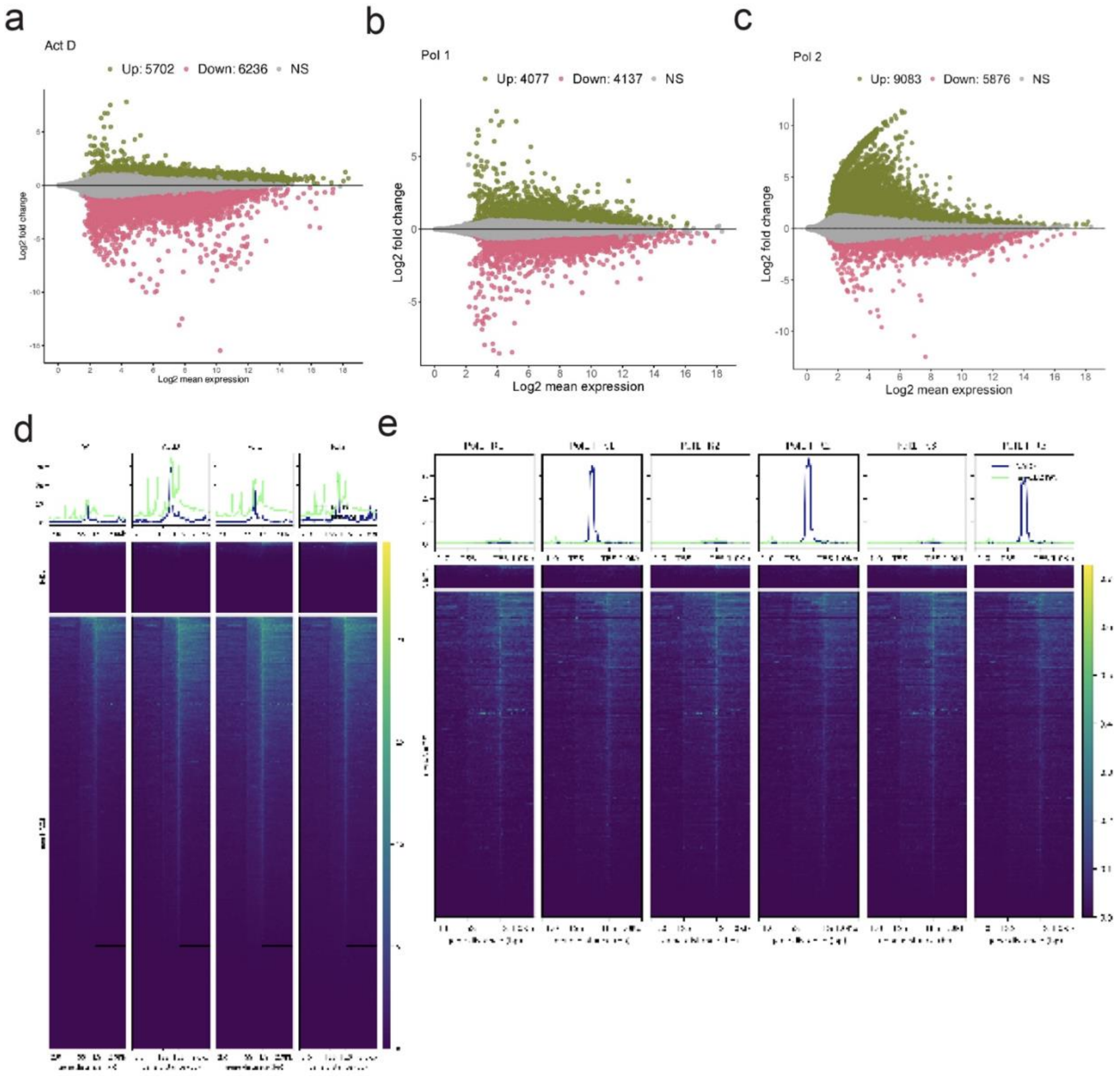

**SI Figure 4.** Additional RNA-seq analysis. Degron lines (POLR2A-AID2 or POLR1A-AID1) were treated with auxin for 8 hours, DMSO for 8 hours, or Actinomycin D treated (5 µg/mL) for 6 hours prior to transcriptomic sequencing. (**A-C**) MA plots for each condition showing log2 expression as a function of log fold change. (**D**) Coverage plots over gene bodies for each condition in and outside of NADs. (**E**) Coverage plots over gene bodies of differentially expressed genes following degradation of POLR1A in and outside of NADs using.

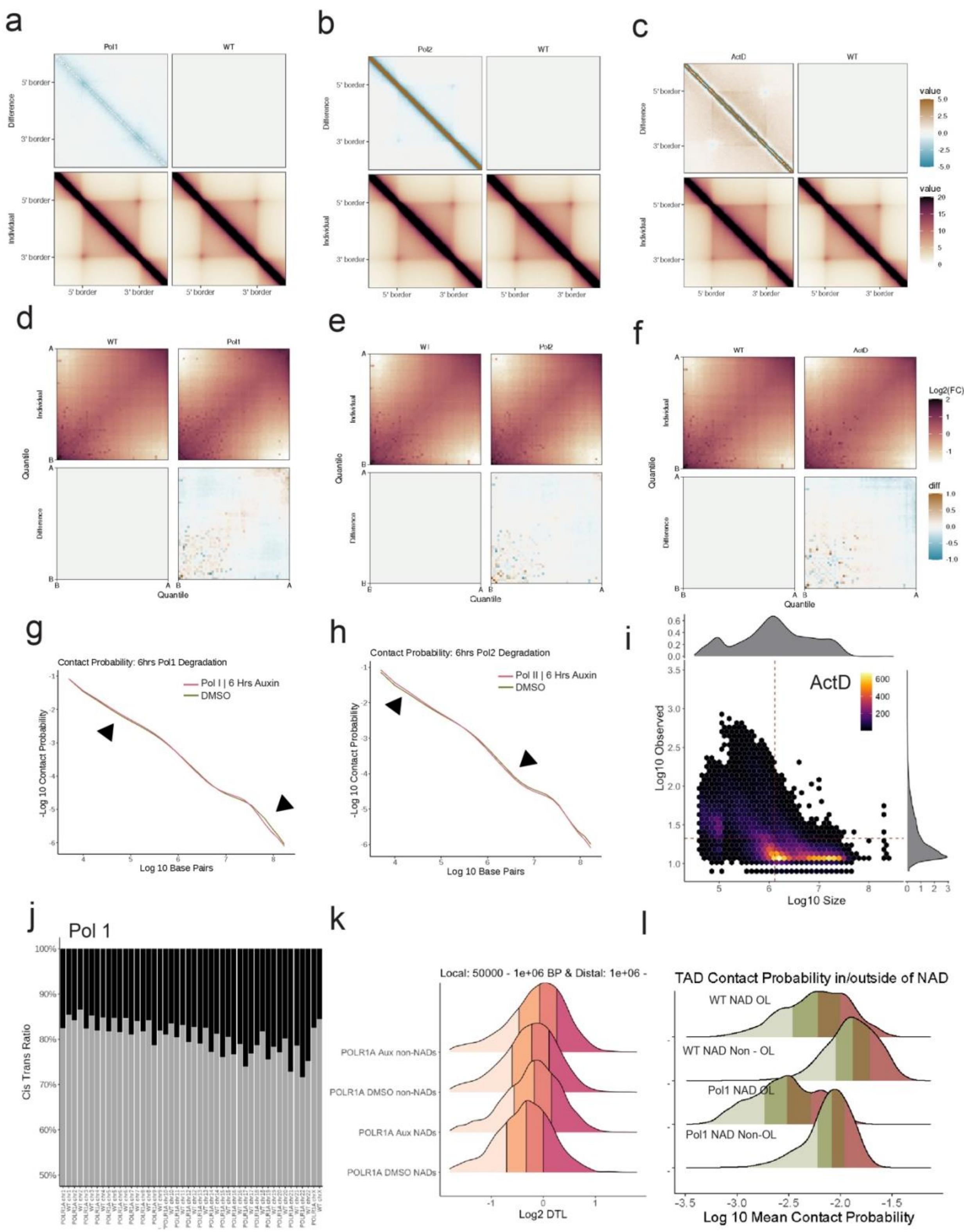

a
Pol1
WT
Difference
Individual
5' border
3' border
b
Pol2
WT
c
ActD
WT
value
d
WT
Pol1
Quantile
Individual
Difference
e
Pol2
f
ActD
Log2(FC)
diff
g
Contact Probability: 6hrs Pol1 Degradation
Pol I | 6 Hrs Auxin
DMSO
-Log 10 Contact Probability
Log 10 Base Pairs
h
Contact Probability: 6hrs Pol2 Degradation
Pol II | 6 Hrs Auxin
DMSO
i
ActD
Log10 Observed
Log10 Size
j
Pol 1
Cis Trans Ratio
k
Local: 50000 - 1e+06 BP & Distal: 1e+06 -
POLR1A Aux non-NADs
POLR1A DMSO non-NADs
POLR1A Aux NADs
POLR1A DMSO NADs
Log2 DTL
l
TAD Contact Probability in/outside of NAD
WT NAD OL
WT NAD Non - OL
Pol1 NAD OL
Pol1 NAD Non-OL
Log 10 Mean Contact Probability

**SI Figure 5.** Additional Hi-C analysis. Degron lines (POLR2A-AID2 or POLR1A-AID1) were treated with auxin for 6 hours, DMSO for 6 hours, or Actinomycin D treated (5 μg/mL) for 1 hour prior to contact mapping. For untreated HCT116 cells and treated degron lines: (**A-C**) TAD insulation plots (GENOVA), (**D-E**) compartment pile-up plot (GENOVA), and (**G-I**) Relative contact probability. (**J**) Cis/trans rate per chromosome for POLR1A-degraded cells. (**K**) Distribution of Distal-to-Local ratio values calculated in 10 kb bins for POLR1A-degraded cells in and outside of NADs. (**I**) Scatterplot of log10 loop strength against log10 size for each loop in ActD treated cells. Loops called using HICCUPS. Heatmap shows loop density.

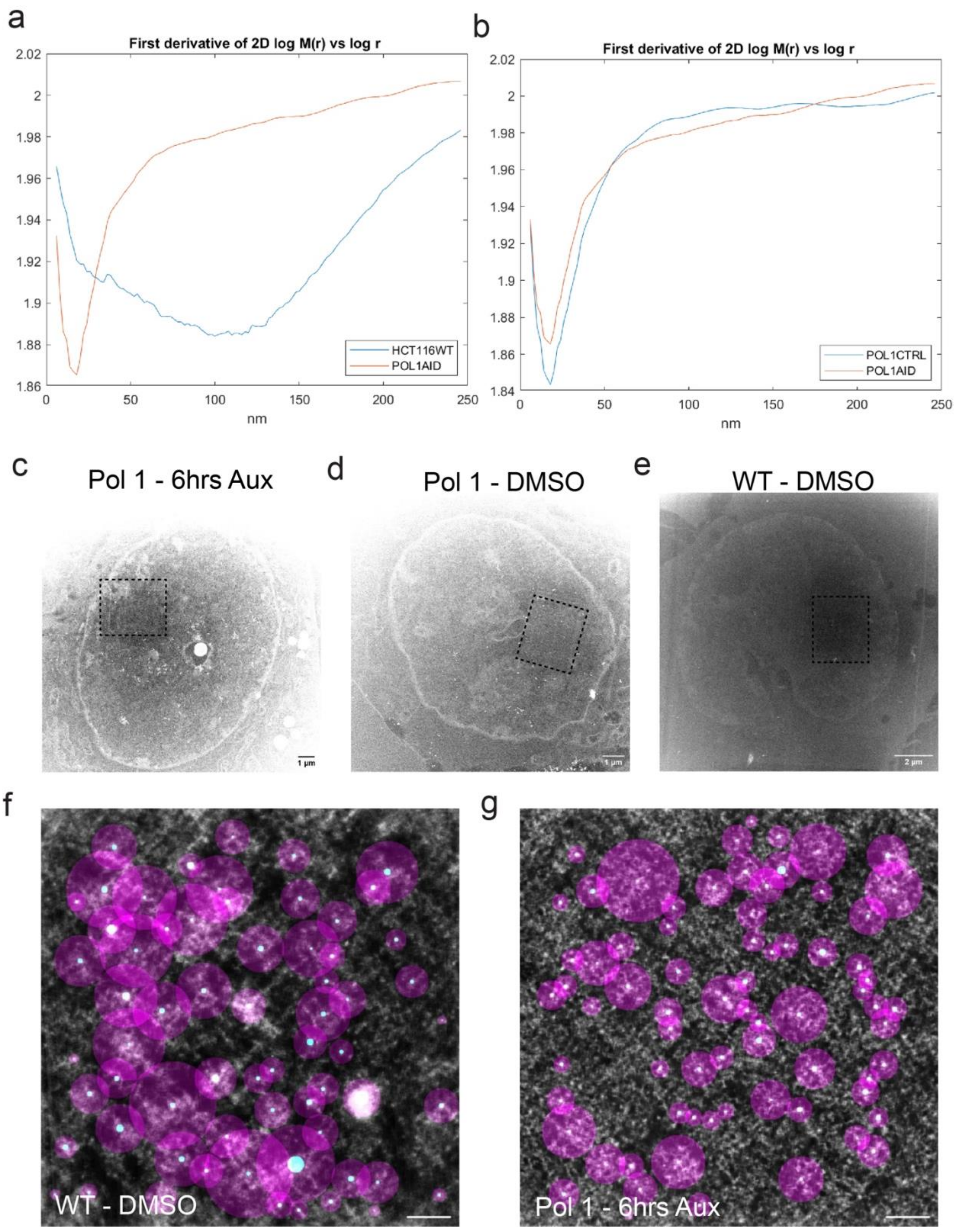

a
First derivative of 2D log M(r) vs log r
2.02
2
1.98
1.96
1.94
1.92
1.9
1.88
1.86
0
50
100
150
200
250
nm
HCT116WT
POL1AID
b
First derivative of 2D log M(r) vs log r
2.02
2
1.98
1.96
1.94
1.92
1.9
1.88
1.86
1.84
0
50
100
150
200
250
nm
POL1CTRL
POL1AID
c
Pol 1 - 6hrs Aux
1 μm
d
Pol 1 - DMSO
1 μm
e
WT - DMSO
2 μm
f
WT - DMSO
g
Pol 1 - 6hrs Aux

**SI Figure 6.** Supplementary chromSTEM analysis. (**A-B**) The local derivative of log-log scale cumulative mass vs distance correlation plot log(M) vs log(r), representing the fitting of scaling perimeter D of chromatin polymer. (**C-E**) Scanning Transmission Electron Microscope HAADF image of nucleus. The box shows the region where ChromSTEM tomography data is collected. (**F-G**) ChromSTEM tomogram overlayed with identified domains center (highlighted in yellow) based on local maxima (ImageJ) and domain size in purple used for **Figure 5I**. Scale bar = 200nm.

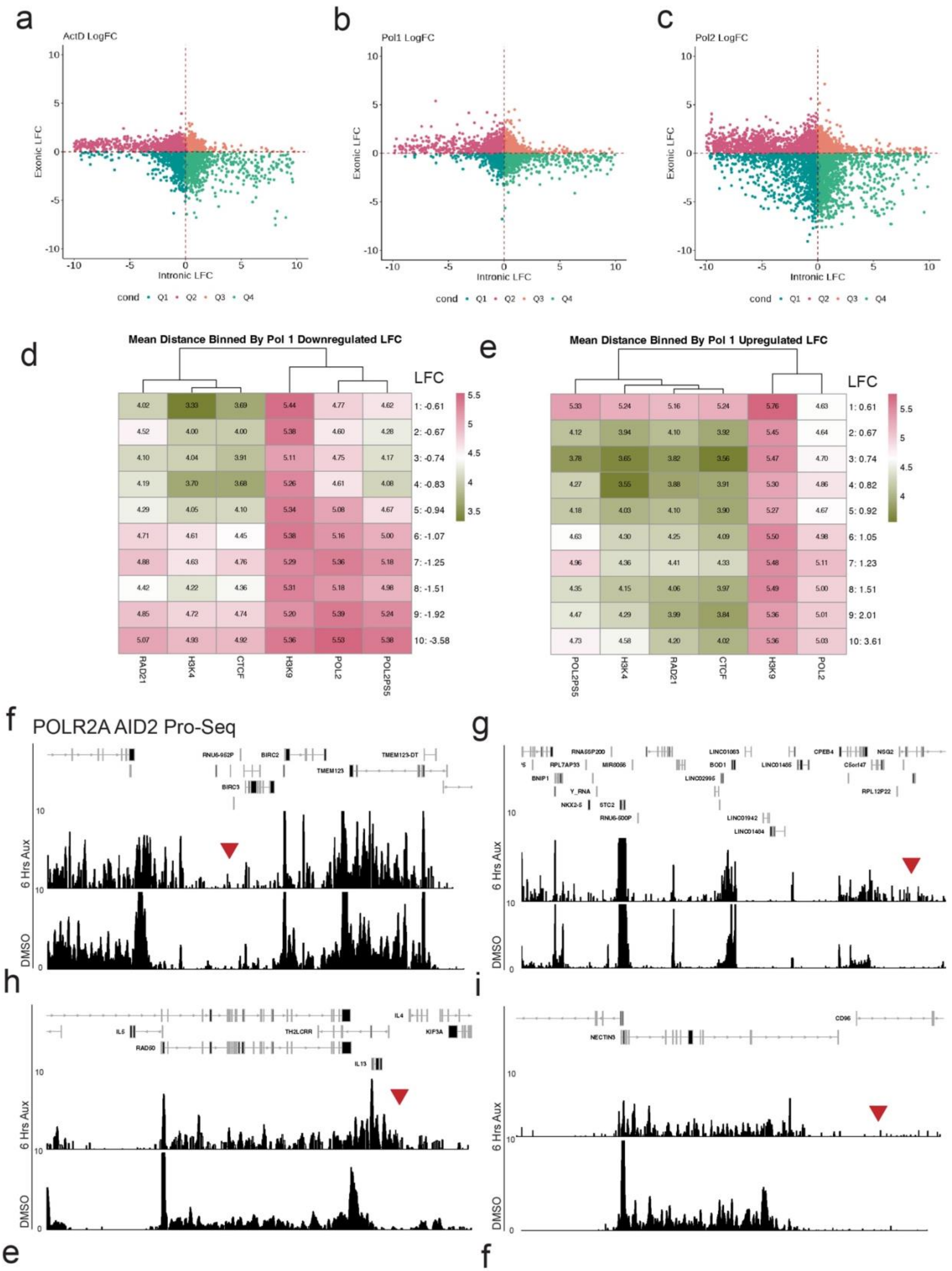

a
ActD LogFC
b
Pol1 LogFC
c
Pol2 LogFC
Exonic LFC
Intronic LFC
cond Q1 Q2 Q3 Q4
d
Mean Distance Binned By Pol 1 Downregulated LFC
RAD21 H3K4 CTCF H3K9 POL2 POL2PS5
LFC
1: -0.61 4.02 3.33 3.69 5.44 4.77 4.62
2: -0.67 4.52 4.00 4.00 5.38 4.60 4.28
3: -0.74 4.10 4.04 3.91 5.11 4.75 4.17
4: -0.83 4.19 3.70 3.68 5.26 4.61 4.08
5: -0.94 4.29 4.05 4.10 5.34 5.08 4.67
6: -1.07 4.71 4.61 4.45 5.38 5.16 5.00
7: -1.25 4.88 4.63 4.76 5.29 5.36 5.18
8: -1.51 4.42 4.22 4.36 5.31 5.18 4.98
9: -1.92 4.85 4.72 4.74 5.20 5.39 5.24
10: -3.58 5.07 4.93 4.92 5.36 5.53 5.38
e
Mean Distance Binned By Pol 1 Upregulated LFC
POL2PS5 H3K4 RAD21 CTCF H3K9 POL2
LFC
1: 0.61 5.33 5.24 5.16 5.24 5.76 4.63
2: 0.67 4.12 3.94 4.10 3.92 5.45 4.64
3: 0.74 3.78 3.65 3.82 3.56 5.47 4.70
4: 0.82 4.27 3.55 3.88 3.91 5.30 4.86
5: 0.92 4.18 4.03 4.10 3.90 5.27 4.67
6: 1.05 4.63 4.30 4.25 4.09 5.50 4.98
7: 1.23 4.96 4.36 4.41 4.33 5.48 5.11
8: 1.51 4.35 4.15 4.06 3.97 5.49 5.00
9: 2.01 4.47 4.29 3.99 3.84 5.36 5.01
10: 3.61 4.73 4.58 4.20 4.02 5.36 5.03
f
POLR2A AID2 Pro-Seq
RNU6-952P BIRC2 TMEM123-DT TMEM123 BIRC3
6 Hrs Aux
DMSO
g
RNA5SP200 LINC01063 CPEB4 NSG2 RPL7AP33 MIR8086 BOD1 LINC01485 C5orf47 BNIP1 LINC02995 Y_RNA RPL12P22 NKX2-5 STC2 RNU6-500P LINC01942 LINC01484
6 Hrs Aux
DMSO
h
IL4 IL5 TH2LCRR KIF3A RAD50 IL13
6 Hrs Aux
DMSO
i
CD96 NECTIN3
6 Hrs Aux
DMSO
e
f

**SI Figure 7.** Supplimentary transcriptomic and ChIP analysis. (**A-C**) Exon log fold change and intron log fold change generated using Index plotted for each gene. (**D**) Heatmap of distance from protein or post-translational modification to nearest downregulated gene in POLR1A-degraded cells, binned by LFC. Color corresponds to log10 distance to mark. (**E**) Same as (**D**) but for nearest upregulated gene. (**A-C**) Publicly available POLR2A-degraded Pro-Seq. Genomic loci correspond to **Figure 7D-G** and red arrows point to same read-through regions.

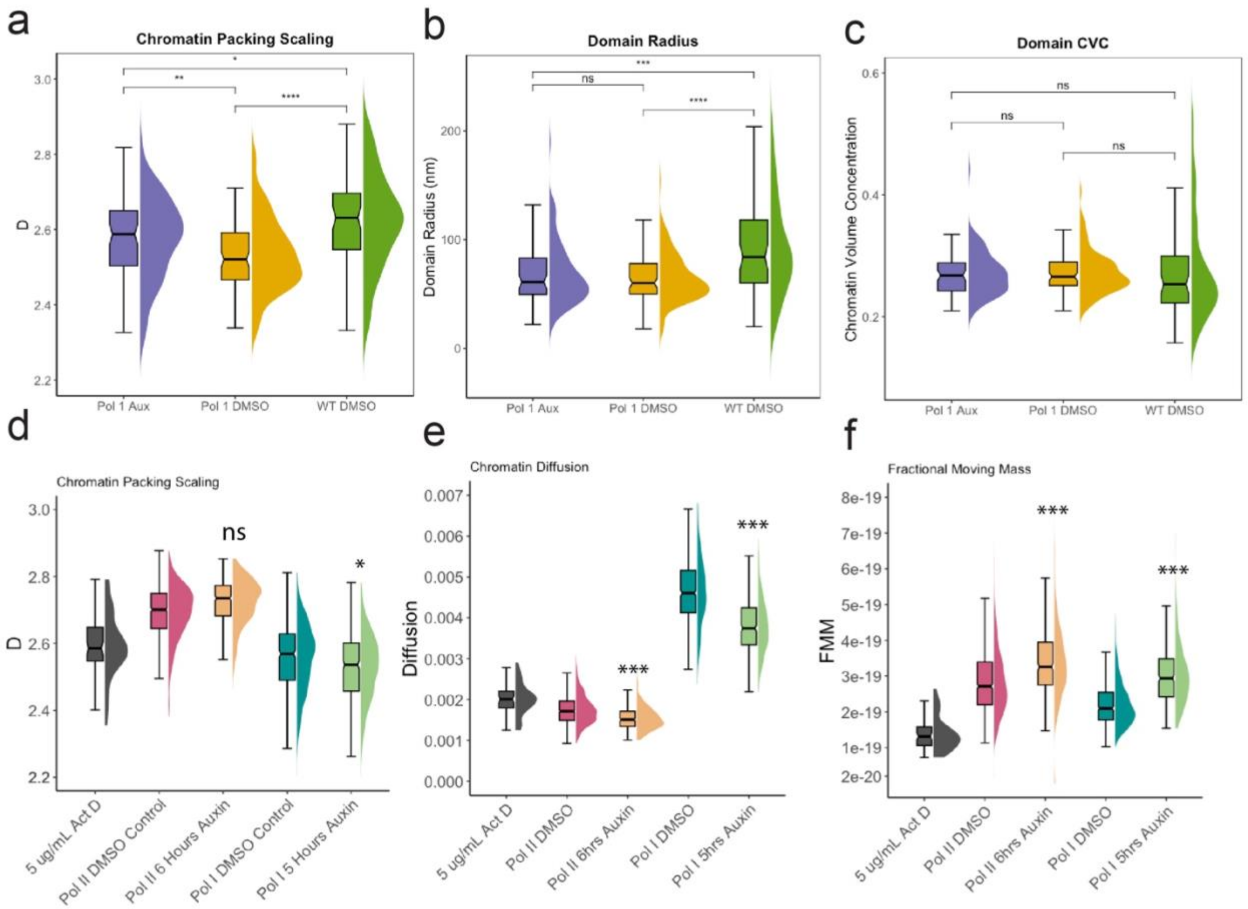


**SI Figure 8.** Additional PWS and ChromSTEM analysis. (**A-C**) Distribution of individual domain properties observed on ChromSTEM from **Figure 5J-L** but including POLR1A-AID1 treated with DMSO for 6 hours: packing scaling, domain radius, and chromatin volume concentration of individual domains, respectively. (**D-F**) PWS following treatment from (**Figure 4I-K**) but including POLR1A-AID1 treated with DMSO for 6 hours: average nuclear packing scaling, diffusion, and fractional moving mass of chromatin domain nuclear average, respectively.

### Table S1. Supplementary Data

| Repo | Experiment | File | Method | Source | Protein or Mark | Type |
|---|---|---|---|---|---|---|
| 4DN | 4DNES3PG7E7F | 4DNFI4HQPGVC | Dam-ID | HCT116 | 4X-AP3D1 (NADs) | BED |
| GEO | GSE150822 | GSE150822_Coordinates_ NADonly_NADsLADs_LADonly.xlsx | Dam-ID | mESC | NoLS (NADs) | BED |
| GEO | GSE145791 | SRR11150184-SRR11150187 | ChIP-Seq | mESC | POLR1A | FASTQ |
| GEO | GSE145791 | SRR11150188-SRR11150191 | ChIP-Seq | mESC | POLR2A | FASTQ |
| GEO | GSE145791 | SRR11150182-SRR11150183 | ChIP-Seq | mESC | Input | FASTQ |
| Encode | ENCSR945WVF | ENCFF768JPY,ENCFF454CLB, ENCFF896KDP, ENCFF561IFD | Pro-Seq | HCT116 | POLR2A | BigWig |
| Encode | ENCSR240PRQ | ENCFF003KHP | ChIP-Seq | HCT116 | CTCF | BED |
| Encode | ENCSR333OPW | ENCFF394IMO. | ChIP-Seq | HCT116 | H3K4me3 | BigWig |
| Encode | ENCSR179BUC | ENCFF572IBD | ChIP-Seq | HCT116 | H3K9me3 | BigWig |
| Encode | ENCSR000BML | ENCFF229YMU | ChIP-Seq | HCT116 | POLR2A-PS5 | BED |
| Encode | ENCSR000BSB | ENCFF391AAM | ChIP-Seq | HCT116 | Rad21 | BED |
| GEO | GSM1544525 | GSM1544525_Pol_I_ChIP-seq_HMEC.bed | ChIP-Seq | HMEC | POLR1A | BED |

### Table S2. Reagents used in ChromSTEM Staining

| Reagent | Formula |
|---|---|
| Washing solution | Hank’s balanced salt solution without calcium and magnesium |
| Fixation solution | 2.5% EM grade glutaraldehyde<br>2% paraformaldehyde<br>2 mM $CaCl_2$<br>0.1 M sodium cacodylate buffer, pH = 7.4 |
| Blocking solution | 10 mM glycine<br>10 mM potassium cyanide<br>0.1 M sodium cacodylate buffer, pH = 7.4 |
| DNA staining solution | 10 µM DRAQ5<br>0.1% SAPONIN<br>0.1 M sodium cacodylate buffer, pH = 7.4 |
| Bathing solution | 2.5 mM 3,3’- diaminobenzidine tetrahydrochloride (DAB)<br>0.1 M sodium cacodylate buffer, pH = 7.4 |
| Reduced osmium staining solution | 2% osmium tetroxide<br>1.5% potassium ferrocyanide<br>2 mM $CaCl_2$<br>0.15 M sodium cacodylate buffer, pH = 7.4 |
| Durcupan™ resin mixture 1 | 10 mL Durcupan ™ ACM single component A, M, epoxy resin<br>10 mL Durcupan ™ ACM single component B, hardener 964<br>0.15 mL Durcupan ™ ACM single component D |

| | |
|---|---|
| Durcupan™ resin mixture 2 | 10 mL Durcupan ™ ACM single component A, M, epoxy resin<br>10 mL Durcupan ™ ACM single component B, hardener 964<br>0.2 mL Durcupan ™ ACM, single component C, accelerator 960<br>0.15 mL Durcupan ™ ACM single component D |
| 1:1 infiltration mixture | 10 mL 100% ethanol<br>10 mL Durcupan ™ resin mixture 1 |
| 2:1 infiltration mixture | 5 mL 100% ethanol<br>10 mL Durcupan ™ resin mixture 1 |

**Table S3. Antibodies used in this study**

| Source | AB | Assay |
|---|---|---|
| Abcam, ab176916 | rabbit anti-H3K9me3 | STORM/SMLM |
| Abcam, AB252855 | rat anti-RNA Polymerase II | STORM/SMLM/IF |
| Santa Cruz, sc-48385 | Mouse monoclonal anti-RPA194 | STORM/SMLM/IF/CUT&TAG |
| Abcam, ab150167 | Goat Anti-Rat IgG Alexa Fluor 647 | STORM/SMLM/IF |
| ThermoFisher | goat anti-rabbit AF568 | STORM/SMLM/IF |
| ThermoFisher | goat anti-rat AF488 | STORM/SMLM |
| Thermo Fisher , A31572 | Donkey Anti-Mouse IgG H+L Alexa Fluor 647 | IF |
| Santa Cruz, sc-13057 | mouse monoclonal C23 (Nucleolin) | IF |
| Edward Chen, U of F | Human UBF | IF |
| ThermoFisher, A11032 | Goat anti-Mouse IgG (H+L) Alexa Fluo 594 | IF |
| Thermo Fisher, A11008 | Goat anti-Rabbit IgG (H+L) Alexa Fluor 488 | IF |
| ThermoFisher, A21445 | Goat anti-Human IgG (H+L) Alexa Fluor 647 | IF |
| Abcam, ab5408 | Anti-RNAPII-PS5 | WB |
| Promega, #W4018 | anti-rabbit IgG HRP | WB |
| Active Motif, 39916 | Rabbit polyclonal Histone H3K4me3 a | CUT&TAG |
| Active Motif, 39065 | Rabbit polyclonal Histone H3K9me3 | CUT&TAG |
| Active Motif, 39034 | Rabbit polyclonal Histone H3K27ac | CUT&TAG |
| Active Motif, 91152 | Abflex Mouse RNA Polymerase II | CUT&TAG |
